# Phylogenetically diverse wild plant species use common biochemical strategies to thrive in the Atacama Desert

**DOI:** 10.1101/2023.08.25.554423

**Authors:** Thomas Dussarrat, Ricardo Nilo-Poyanco, Tomás C. Moyano, Sylvain Prigent, Tim L. Jeffers, Francisca P. Díaz, Guillaume Decros, Lauren Audi, Veronica M Sondervan, Bingran Shen, Viviana Araus, Dominique Rolin, Dennis Shasha, Gloria M. Coruzzi, Yves Gibon, Claudio Latorre, Pierre Pétriacq, Rodrigo A. Gutiérrez

## Abstract

The best ideotypes are under mounting pressure due to increased aridity in many parts of the world. Understanding the conserved molecular mechanisms that evolve in wild plant species adapted to harsh environments is crucial in developing new strategies for sustainable agriculture. Yet our knowledge of such mechanisms in wild species is scant, particularly in extreme environments. We performed metabolic pathway reconstruction using transcriptome information from 32 Atacama plant species and phylogenetically related plant species that do not live in Atacama (Sister species). We analyzed pathway and reaction enrichment to understand the biochemical commonalities and differences of wild Atacama plant species. To gain insights into the mechanisms that ensure plant survival, we compared expressed gene isoform numbers and gene expression patterns between the annotated biochemical reactions from 32 Atacama and Sister species. We found significant biochemical convergences in primary and secondary metabolism characterized by reactions enriched in at least 50% of the Atacama species across major plant phylogenetic lineages. Analysis of the annotation indicated potential advantages against drought, salinity, high solar irradiance, and nitrogen starvation. These findings suggest that the adaptation in the Atacama Desert may result in part from shared genetic legacies governing the expression of key metabolic pathways to face harsh environmental conditions. Enriched reactions corresponded to ubiquitous compounds common to extreme and agronomic species and were congruent with our previous metabolomic analyses in these Atacama species. Hence, genes underlying these adaptive traits offer promising candidates for improving abiotic stress resilience in crop species.

## Introduction

Plants are sessile organisms which rely on the availability of local resources to live. Substantial efforts in plant breeding programs have focused on developing increasingly productive and resistant ideotypes to environmental constraints (Voss-Fels et al., 2019). However, accelerated climate change is rendering these genetic improvements insufficient (Bailey-Serres et al., 2019). A few wild plant species flourish under harsh environmental conditions, representing a unique reservoir of adaptive mechanisms and genetic resources (Díaz et al., 2019; Carrasco-Puga et al., 2021). Random genetic mutations have tailored the plant genome to extreme ecosystems such as deserts (Bolger et al., 2014). Understanding plant survival strategies from extreme environments could help unravel resistance mechanisms that would greatly benefit global food security. Although promising, studies on extreme wild plant species are scant and mostly performed under controlled laboratory conditions. Those settings lack ecological context and may hinder the identification of adaptive traits that are expressed only under natural conditions (Dussarrat et al., 2021).

Plants evolved temporal and spatial strategies to optimize the balance between development and defense in hostile ecosystems (Dussarrat et al., 2021). For instance, the survival of plants that cannot adapt in time depends on genetic mechanisms underlying, for example, metabolic processes such as osmoregulation or precise management of the lipidic profile to avoid water loss (Bolger et al., 2014; Turner, 2018). However, most of this knowledge is based on studies of one or a limited number of species. Exploring conserved and shared adaptive mechanisms may increase the likelihood of success when transferred to other species (Turner, 2018). For instance, recent analyses of several tens of thousands of plant species indicated that the appearance of several phenotypic traits strongly correlated with environmental variations (*e.g.* latitudinal gradient) worldwide (Joswig et al., 2021). Therefore, it is of interest to determine the extent to which convergent evolution occurred between molecular mechanisms. We have evidence that this can happen. A recent study from our group unveiled 265 positively selected genes (PSGs) from 32 species from the Atacama Desert, the driest non-polar desert on Earth (Eshel et al., 2021). These genes encompassed various molecular processes related to the protection against high solar irradiance, nitrogen starvation and osmotic stress, with a great part of these PSGs being shared among different plant lineages. Hence, the exciting possibility of a strong influence of shared adaptive genetic processes in adaptation and an intriguing over-expression of genes related to protective metabolite synthesis emerged (Eshel et al., 2021). These results suggest the existence of common selected mechanisms managing key metabolic processes that govern plant life under major abiotic threats. However, the exact nature, role, and evolutionary trajectories of the enriched biochemical reactions and pathways in Atacama plant species remain unclear.

Studies demonstrated that plant metabolism is an excellent predictor of environmental variation in harsh biomes (Fiehn, 2002; Kumari et al., 2020). For instance, the orchestration of primary and secondary metabolism led to the accumulation of amino acids, phenolics, and nitrogen-containing metabolites (Lugan et al., 2010; Dussarrat et al., 2021). While few studies performed an ecological metabolomic approach using multiple species, promising results demonstrated a significant correlation between phytochemical diversity and environmental variation (Defossez et al., 2021). Recent work used 24 Atacama plant species to discover a metabolic toolbox composed of 39 metabolites predicting plant environment independently of the plant lineage. These common predictors were also detected in agronomic plant species, raising great hope for their use in engineering crop resilience to harsh abiotic constraints (Dussarrat et al., 2022). However, the genetic traits governing the modulation of this generic toolbox remain unknown.

The Atacama Desert is an extreme and challenging environment where the intensity of abiotic stresses shapes plant life (Eshel et al., 2021). The Atacama Desert is currently characterized by extremely low precipitations (less than 20 to 160 mm/year) and high solar irradiance (600 W/m²/d) compared to other deserts or high mountain ecosystems (Báez and Collins, 2008; Zhang et al., 2010; Jordan et al., 2014; Díaz et al., 2016; Ziaco et al., 2018). In addition, plants face extremely low levels of macronutrients such as nitrogen and high levels of salinity (Eshel et al., 2021). Thus, the Atacama Desert offers unique opportunities to uncover molecular mechanisms determining plant performance under extreme conditions.

This study aims to decipher the adaptive biochemical responses selected through the evolution of multiple plant lineages from the Atacama Desert. We used existing computer pipelines to study the evolutionary trajectories of plant biochemical compounds using genome-wide sequencing data (Chae et al., 2014; Schläpfer et al., 2017; Kang et al., 2020). Have evolutionary processes led to the convergent fixation of various biochemical reactions and pathways or conversely resulted in strategy diversification? To answer this question, we first extracted biochemical reaction-related genes and annotated associated pathways using transcriptome data of 32 Atacama species covering fourteen plant families. These 32 Atacama species are the most relevant species found in the hyperarid core of the Atacama Desert based on their coverage, persistence and distribution (Eshel et al., 2021). We performed a similar workflow in 32 Sister species that were phylogenetically related to the Atacama species but lived in other milder environments (Eshel et al., 2021). We analyzed the number of gene expressed gene isoforms per reaction and qualitative gene expression levels, and evaluated the enrichment of biochemical reactions and pathways when comparing each pair of Atacama and related species using over-representation analysis (ORA) (Wieder et al., 2021). This computational strategy highlighted the convergent selective advantages of Atacama plant metabolomes. The most ubiquitous genetically enriched responses were related to protective mechanisms against major abiotic stresses in the Atacama Desert, such as drought, nitrogen deprivation and high light intensity. These findings provide new insights into adaptive mechanisms for plant survival in the Atacama Desert and new genetic targets for crop engineering for the sake of more resilient agriculture.

## Materials and methods

### Plant material

Reaction and pathway enrichment analyses were performed using previously described and available transcriptome data from 32 Atacama plant species (Talabre-Lejía transect, lat 22°-24°S) (Eshel et al., 2021). The high quality and completeness of the transcriptome data were validated previously (Eshel et al., 2021). This natural environment spans an altitudinal cline from 2,470 to 4,470 m a.s.l. and involves three vegetation belts defined based on plant species composition (Carrasco-Puga et al., 2021): the Prepuna (low elevation, high salinity and low water availability), the Puna shrubland and the Steppe (high elevation, low temperatures) (Eshel et al., 2021 and Fig. 1). The species present in these vegetation belts covered 14 distinct plant families across the altitudinal cline (Eshel et al., 2021). Transcriptome data from 32 phylogenetically-related species available from public data sources were also used (defined in Eshel *et al*., 2021). Herein, those species are referred to as “Sister species”.

**Fig. 1.**
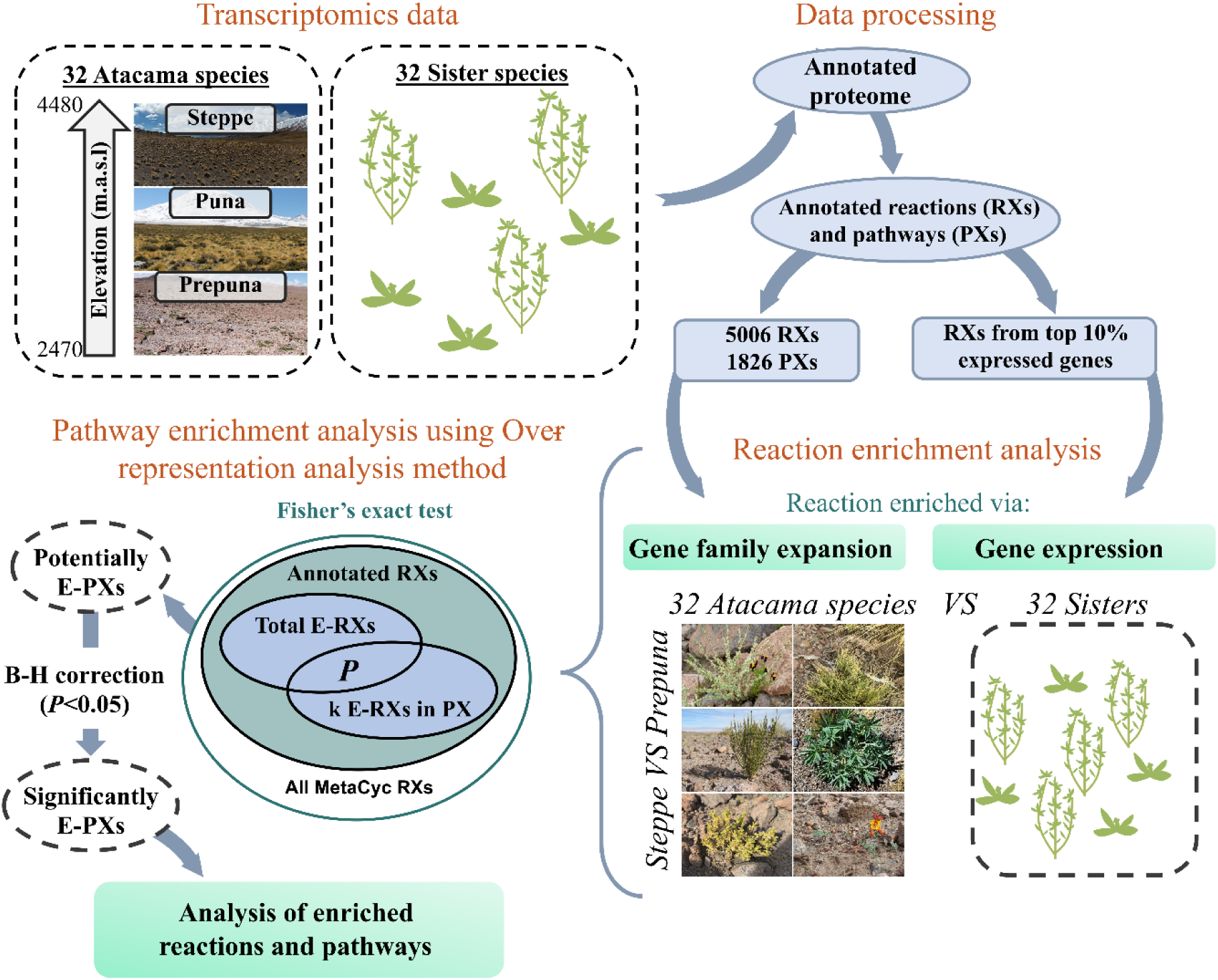
A simplified scheme of the reaction and pathway enrichment approach. *P* represents the probability of finding at least k reactions in pathway X (k E-RXs in PX) and was calculated using Fisher’s exact test based on the hypergeometric distribution (Wieder et al., 2021). *E-RXs*: enriched reactions, *E-PXs*: enriched pathways.

### Generation of annotated reactions and pathways using PathwayTools

Reactions and pathways for the 64 species were built using the sequencing data and the e2p2v4 enzyme annotation tool (Schläpfer et al., 2017). Next, PTOOLS v24.5 was used to infer reactions and pathways using default parameter values (Karp et al., 2021).

### Data treatment

We constructed two data sets that were used for further analyses. First, we identified the number of expressed gene isoforms in the Atacama plant species transcriptome data set. To get these numbers, we counted the total number of genes per reaction for each annotated reaction. This analysis provided a first matrix which served as the basis for the reaction enrichment analysis (Fig. 1, Tab. S1). Second, we identified highly expressed genes in Atacama species. To do this, we took the top 10% expressed genes from each of the 32 Atacama plant species transcriptome, as well as the top expressed genes from the 17 related Sister species for which raw sequencing data were available (Eshel et al., 2021). Top expressed genes that were not associated with any reaction were removed. We then defined the number of top-expressed genes per reaction. This number can range from 0 to “*n*” (no or “*n*” top-expressed genes linked to the reaction *i*). This process produced the second matrix, which was used to analyze overexpressed reactions in Atacama based on qualitative gene expression levels (Fig. 1, Tab. S2).

### Reaction enrichment analysis

Reaction enrichment analysis was performed using the two data matrices produced in the previous section: the number of expressed gene isoforms or qualitative expression levels for genes that were annotated to enzymatic reactions. First, exclusive reactions from Atacama or Sister species were extracted (Tab. S3). Second, common reactions (detected in at least one species in each set) were identified. We compared the number of expressed gene isoforms annotated to a reaction in any given Atacama species to the corresponding number of expressed gene isoforms annotated to the same reaction in the Sister species. We did this comparison using the entire data set or partitioned by vegetation belt (*i.e.* Prepuna, Puna, Steppe). Reactions were considered more diverse (*i.e.* expanded) if there were at least 3 times more expressed gene isoforms per reaction when comparing Atacama *versus* Sister species. This threshold was selected because the number of expanded reactions seemed to reach a plateau at this 3-fold ratio for all comparisons made (*i.e.* increasing the ratio did not substantially decrease the number of enriched reactions) (Fig. S1). Since this approach highlighted strong variations in enriched reaction patterns between Prepuna and Steppe, pairwise comparisons between the 32 Atacama-Sister species pairings were performed to avoid the dilution effect. Results were then aggregated to calculate a percentage of occurrence per set (Atacama or Sister) or vegetation belt (Prepuna or Steppe) (Tab. S4).

A second approach focused on gene expression levels by comparing the number of top-expressed gene isoforms per reaction. Average (average in all Atacama species or Prepuna or Steppe environments *versus* average in related Sisters) and pairwise comparisons were performed for each available Atacama-Sister couple. Reactions were considered overexpressed if at least 3 times more associated top-expressed genes were observed in the top 10% of one Atacama species than its related Sister (Fig. S1). Results were aggregated to calculate the percentage of enrichment (Tab. S5).

### Pathway enrichment analysis

To evaluate whether specific pathways were enriched in Atacama plant species (Tab. S6), we calculated the probability *P* of finding at least k reactions in pathway *i* using Fisher’s exact test and the hypergeometric distribution. All analyses were performed in R (version 4.0.4) using existing functions (*e.g.* phyper function) and custom-made scripts (R Core Team, 2021; Wieder et al., 2021) (Fig. 1). *P*-values were adjusted using Benjamini-Hochberg method on enriched pathways, and a corrected *P*<0.05 cutoff was used to define enrichment (Benjamini and Hochberg, 1995) (Tab. S7 and S8). Finally, reactions and pathway enrichment analyses were performed using this same process to compare Steppe and Prepuna evolution. For each species, the number of expressed gene isoforms per reaction and the number of top-expressed gene isoforms per reaction were compared to the average of the opposite vegetation belt. Reactions were considered more diverse or overexpressed if there were at least 3 times more expressed gene isoforms per reaction or 3 times more top-expressed gene isoforms per reaction when comparing a given species *versus* the average of all species from the opposite vegetation belt. The percentage of occurrence per ecosystem was then calculated (Tab. S9). Finally, pathway enrichment analysis was conducted using Fisher’s exact test and the hypergeometric distribution (Tab. S10).

### Annotation of the enriched reactions and pathways

Reactions expanded or overexpressed in at least 50% of the plant species per vegetation belt (*i.e.* at least 50% of the species from Steppe, Prepuna or all Atacama plants when compared to related Sisters) were annotated using the MetaCyc database (Caspi et al., 2020). Besides, metabolism, sub-pathways and biochemical pathways were determined using KEGG and HMDB databases (Kanehisa et al., 2014; Wishart et al., 2018).

### Testing for positively selected genes (PSG) within vegetation belts

Due to reactions enriched in specific vegetation belts, we ran PSG analysis to unveil proteins positively selected for the unique environments of Atacama vegetation belts, an addition to the published PSGs of Atacama *vs* Sisters (Eshel et al., 2021) (Tab. S11 and S12). We processed previous transcriptomics data through the previously established PhyloGenious pipeline (Eshel et al., 2021) to determine the ortholog groups and their best parsimonious tree structure for 32 Atacama transcriptomes and 6 reference transcriptomes (*Arabidopsis thaliana*, *Brachypodium distachyon*, *Glycine max, Physcomitrium patens*, *Solanum lycopersicum*, and *Zea mays*). To find PSGs adapted to Prepuna or Steppe environments, two methods from the HyPhy package (v.2.5.24) were implemented to cross-validate positively selected orthologs groups associated with adaptation to a vegetation belt (Pond et al., 2005; Murrell et al., 2015; Smith et al., 2015). First, BUSTED (v.2.0) tested a collection of branches of a test-group (*i.e.* plants from a specific vegetation belt) against background branches (*i.e.* reference species + all other vegetation belts). Orthologous groups with a minimum of 4 branches were tested. We required that BUSTED results meet the criteria that dN/dS >1 (LRT test, *P* < 0.05) for the test-group branches and dN/dS ≤1 for background branches. The same ortholog groups had to also pass aBSREL (v.1.2) analysis requirement that an individual tested branch had significant dN/dS >1 while no background branches had dN/dS >1. Additionally, PSOGs that passed aBSREL but not BUSTED (dN/dS >1, *P* < 0.05) were kept. These two independent methods ensured that selection PSGs occurred in more than one species, was specific to a vegetation belt, and was detectable by distinct algorithmic methods (Tab. S12).

### Modeling of in-field metabolomics data to enriched reactions

To assess the relevance of enriched reactions and pathways at the metabolic level, we tested the link between enriched reactions and a generic metabolic toolbox predicting plant environment in the same Atacama gradient (Dussarrat et al., 2022) (Tab. S13). The mean normalized intensity of the 39 compounds predicting plant environment (see Dussarrat *et al.,* 2022 for normalization) was calculated per species in R (v. 4.1.1) (R Core Team, 2021). Pearson correlation of each mean metabolite distribution across species was conducted against the number of expressed gene isoforms per reaction or the number of top 10% expressed gene isoforms per reaction with the base R *cor* function using default settings. When, a metabolite was detected in both positive and negative ionization mode or was represented by multiple fragments (*e.g.* N1,N5,N10-Tricoumaroyl spermidine), the best reaction to spectra ID correlation was chosen. A reaction was considered correlated to a metabolite if Pearson r ≥ 0.7 and *P* < 0.05. We additionally noted how often a correlated reaction was Atacama versus Sister enriched, either by isoform number or expression enrichment. This included the highly stringent convergent enrichment requirement of 50% of relevant species, as well as a less stringent requirement that reactions should be enriched in 3 or more Atacama species than in Sisters (Tab. S13 and S14). The tree of 16 species with metabolomics and transcriptome annotation was adapted from the total evidence species tree created by Eshel *et al.,* 2021 using the *keep.tip* function from the *ape* package (v. 5.7) to retain species with metabolomics data and was displayed using the R package *ggtree* (v. 3.6) (Paradis and Schliep, 2019; Yu, 2022).

Separately, a machine-learning model comparing expressed gene isoforms per reaction to mean metabolite concentrate additionally ranked the most informative reactions underlying a given metabolite distribution. An XGBoost regressor (Chen and Guestrin, 2016) ranked feature importance of reaction number per species to predict a given metabolite mean concentration, followed by a 10-fold cross validation process. In each of the 10 runs, 90% of the reaction ID numbers to metabolite concentration relationships were used as a training set, to test 10% of the remaining metabolites. The average validation score of the model and the average feature importance metrics for each reaction was then calculated. The model score ranged from negative infinity to 1, where 1 represents a perfect prediction in validation while 0 is a prediction no greater than random prediction using training targets. The importance of a feature is computed as the normalized total reduction of the criterion brought by that feature in all the decision trees used in the regression model, ranging from 0 to 1. The model score and the two most important reactions for each metabolite are listed in Tab. S14.

## Results

### Atacama-exclusive reactions undergo species and environment specificity

As a first step towards understanding the metabolic capacity expressed by Atacama plant species, we annotated and compared reactions and pathways in 32 Atacama plant species to their related Sister species. Reactions and pathways were defined using previously acquired sequencing data from 32 Atacama species and publicly available transcriptomic data from their 32 Sisters (Eshel et al., 2021). A total of 5,006 annotated reactions (Fig. 1) were extracted from the transcriptome data available in the 64 species using PathwayTools. To assess the presence of chemical mechanisms unique to Atacama plants, we categorized this set of annotated reactions into exclusive or common reactions. A set of 4,260 reactions were shared, while 463 were exclusive to the Atacama species data set. These exclusive reactions were likely to be species-specific (Fig. 2). To question the influence of environmental pressures on the chemical strategies employed by Atacama plants, we classified exclusive reactions from the two extreme vegetation belts (*i.e.* Steppe and Prepuna) into biochemical pathways. Exclusive reactions were equally divided between primary and secondary metabolisms in Prepuna, while 76% represented secondary pathways in Steppe species (Fig. 2). We found fatty acyls, carbohydrates, amino acids and monoterpenes pathways in the exclusive data set in Prepuna species, which are exposed to high osmotic pressure. Conversely, carotenoids and flavonoids were the main pathways in Steppe, an ecosystem characterized by low temperatures, for instance (Fig. 2, Tab. S3).

**Fig. 2.**
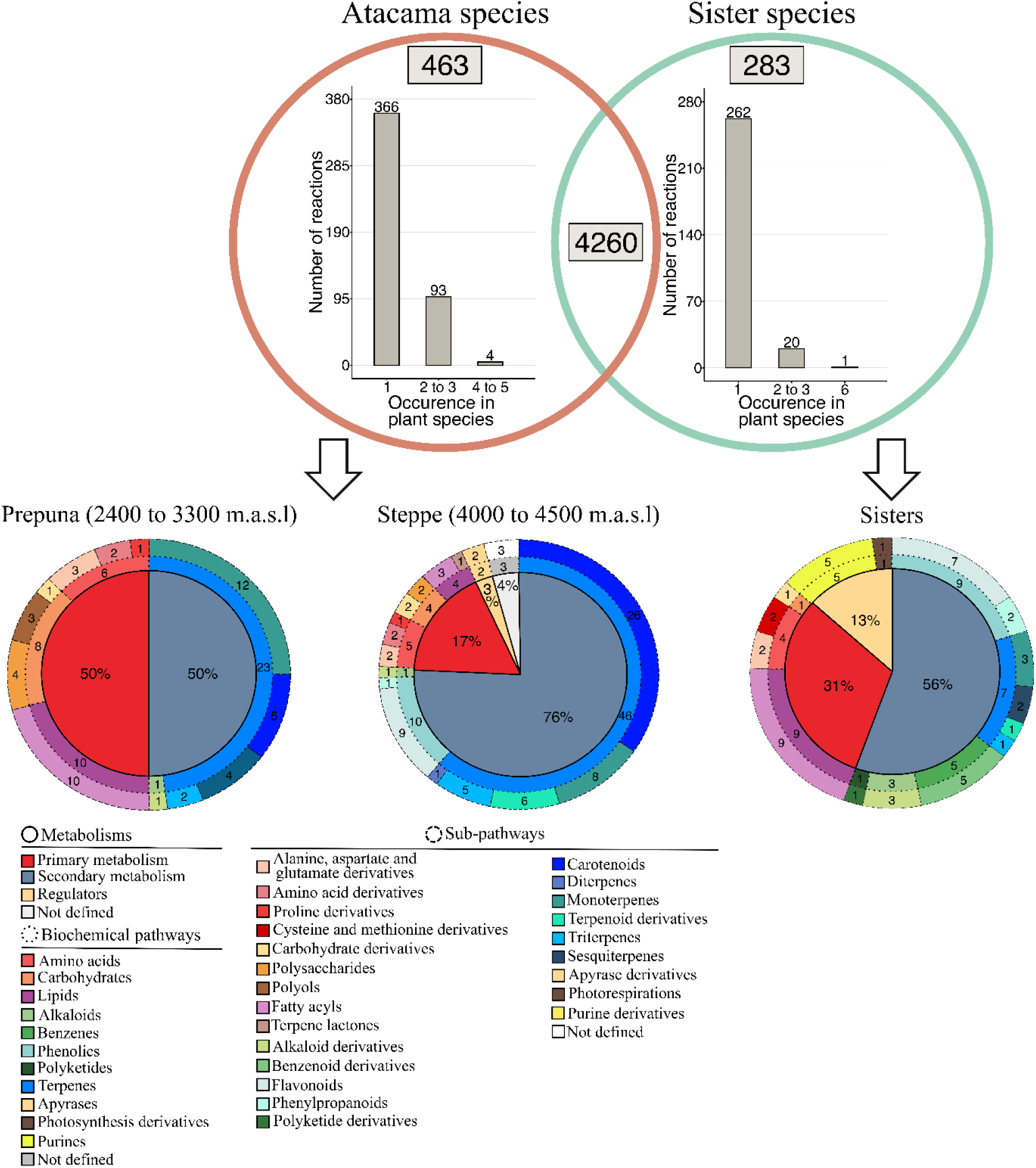
Analysis of the exclusive reactions from Atacama and Sister species. The Venn diagram describes the ecosystem-specific reactions from the 32 Atacama plant species or their closely related Sisters. Metabolism, biochemical pathways and sub-pathways were defined based on MetaCyc and KEGG data. *m.a.s.l*: meters above sea level.

### Shared reactions undergo variations in gene isoform expansion (*i.e.* diversity) or expression

Next, we tested diversity in gene isoforms and gene expression patterns among the genes that code for the 4,260 shared reactions. To evaluate the genetic enrichment, we performed a comparison of expressed gene isoforms and gene expression levels (*i.e.* expressed genes per reaction and top-expressed genes per reaction, respectively) between (i) all Atacama plants *versus* all related Sisters or between (ii) all Steppe or Prepuna species *versus* all related Sisters. Reactions were considered expanded if the number of expressed gene isoforms per reaction was at least 3 times higher in Atacama species as compared to related Sister species (Fig. S1). A 3-fold cutoff was selected because the number of expanded reactions reached a plateau at this ratio. While some reactions were expanded in all vegetation belts, others showed environment specificity (Fig. S2). We subsequently performed pairwise comparisons taking one Atacama *versus* one Sister species to avoid a potential dilution effect due to the environment. These 32 pairwise comparisons yielded a set of 2,507 expanded reactions in at least one Atacama species (Tab. S4). Expanded reactions differed in their specificity at both plant species and environmental levels (Fig. S3). Half (*i.e.* 1,002) of the expanded reactions were specific to one or two vegetation belts, while the remaining were spread in the entire biome (Fig. 3a). These results indicated that the extreme conditions have led to dramatic changes in the number of expressed gene isoforms for multiple reactions in various Atacama species.

**Fig. 3.**
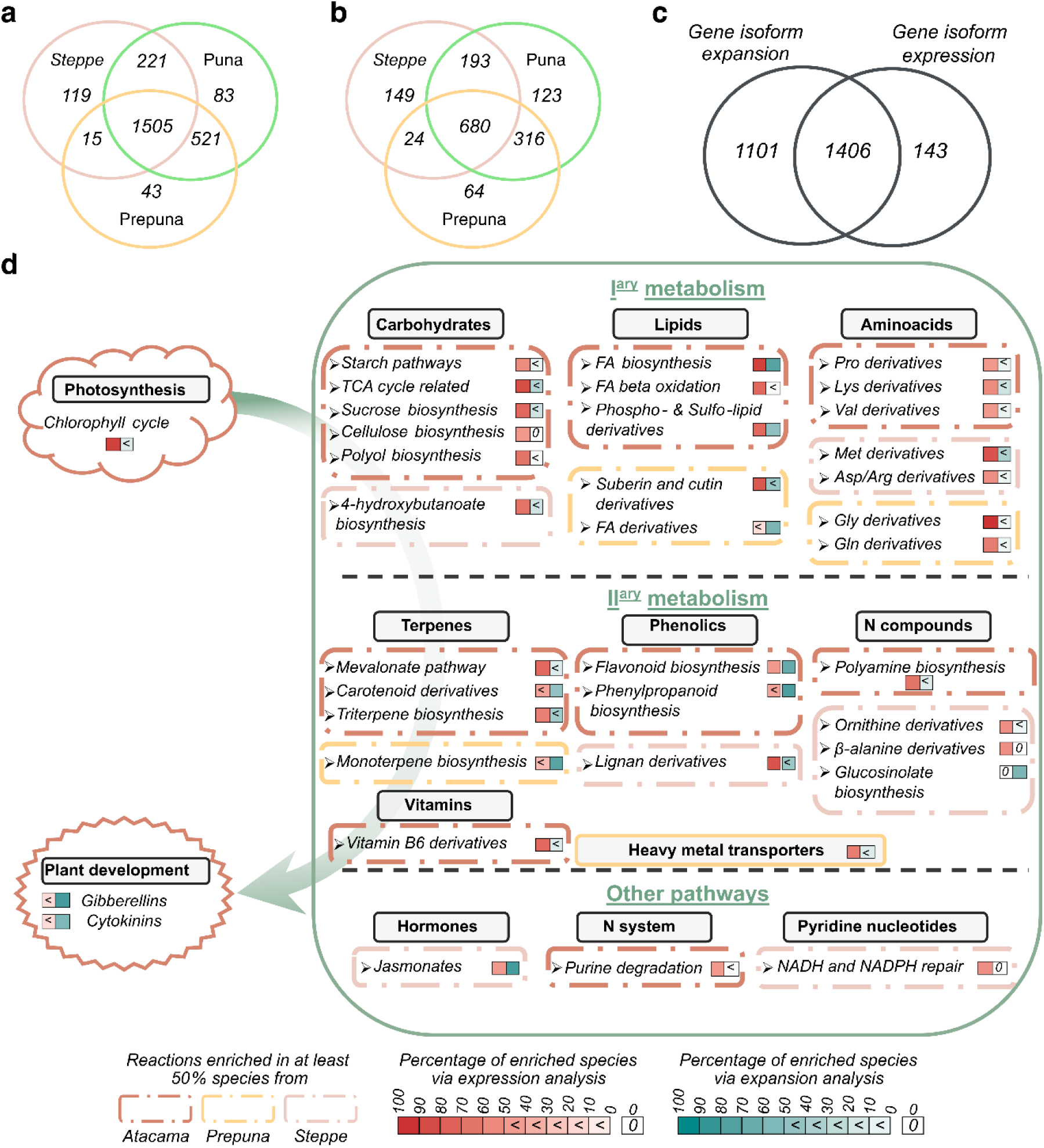
Identification and classification of enriched reactions from Atacama plant species into major biochemical pathways. **a.** Global trends in the distribution of the expanded (*i.e.* diverse) reactions in Atacama species. **b.** Global trends in the distribution of the enriched reactions in Atacama species when comparing the number of top expressed gene isoforms per reaction. **c.** Distribution of diverse and overexpressed reactions in at least one Atacama species. **d.** Results of a pairwise comparison analysis and classification of the reactions expanded and overexpressed in at least 50% of the species. Each Atacama-Sister species pair was analyzed individually and results were then aggregated to calculate a percentage of occurrence per ecosystem (Atacama or Sister) or per vegetation belt (Prepuna or Steppe). *Arg*: arginine*, Asp*: aspartate, *B-H*: Benjamini-Hochberg, *Chl*: chlorophyll, *FA*: fatty acids, *Gln*: glutamine, *Glu*: glutamate, *Gly*: glycine, *Lys*: lysine, *Met*: methionine, *Plip*: phospholipids, *Pro*: proline, *Val*: valine.

While having metabolic potential coded in the genome, this potential must be expressed to be functionally relevant for adaptation. As a first step to understanding gene expression variation in metabolic reactions, we extracted and compared the top 10% expressed genes from Atacama and Sister species (Eshel *et al.,* 2021 and Fig. 1). We asked how many top 10% expressed gene isoforms per specific reaction were included in this set and defined a highly expressed reaction when the number of gene isoforms for this reaction was three times higher in any given comparison: all Atacama *versus* Sister species and pairwise comparisons taking one Atacama versus one sister species (Fig. S1). While gene isoform diversity patterns were analyzed on the 32 couples, 17 couples were used to study gene expression levels since raw sequencing data were only available for 17 Sister species. Pairwise comparisons pinpointed a set of 1,549 highly expressed reactions in Atacama species (Tab. S5, Fig. S3). Results showed that about half (44%) of the highly expressed reactions were observed in all vegetation belts (Fig. 3b). 91% of the highly expressed reactions were associated with a higher number of gene isoforms in Atacama species (Fig. 3c), indicating a very significant overlap between enriched reactions via gene isoform diversity and expression analyses. Overall, these results provided a unique set of diverse and highly expressed reactions in extreme plant species and suggested the existence of conserved metabolic mechanisms in extremophile plant species.

### Enrichment analyses unveil convergent biochemical strategies

To test the hypothesis that Atacama plant evolution led to the convergent enrichment of biochemical reactions among the different plant lineages, we extracted the reactions enriched in at least 50% of the species in a given ecosystem to ensure the presence of various plant lineages. The gene isoform diversity analysis results illustrated the diversity of reactions in at least sixteen species when considering all Atacama plants or at least seven species when considering individual vegetation belts (*i.e.* Prepuna or Steppe) (Fig. 3d). Similarly, results from gene expression analysis encompassed highly expressed reactions in at least nine or four species since raw sequencing data were available for seventeen Sister species. Enriched reactions were shared between species from (i) the entire transect or (ii) specific vegetation belts and suggested a global re-orchestration of primary and secondary metabolism.

### Convergent biochemical strategies shared independently of vegetation belts

Shared reactions (*i.e.* at least 50% of species) related to photosynthesis and primary pathways were mainly revealed by gene expression analysis, while also being more diverse in multiple species (Fig. 3d). A significant proportion of these highly expressed reactions referred to the synthesis of a battery of protective primary compounds. Reactions associated with the regulation of compounds from carbohydrate (*e.g.* starch and polyols), lipid (*e.g.* waxes synthesis) and amino acid pathways (*e.g.* proline) were overexpressed in the majority of Atacama plant species (Fig. 3d). Most of the highlighted reactions by our computational analysis were previously characterized for their function in mitigating various abiotic stresses. For example, glucan/water dikinase, disproportionating enzyme, fatty aldehyde decarbonylase, and various inositol phosphatases were greatly overexpressed in Atacama species (Tab. S5, Yano *et al*., 2005; Lou *et al*., 2007; Jia *et al*., 2019). Lipid metabolism was highlighted by both gene isoform diversity and expression analyses (*e.g.* fatty acyl-ACP thioesterases). Diverse reactions pertained to the regulation of hormones such as gibberellins (*e.g.* gibberellin oxidase) and cytokinins (*e.g.* cytokinin oxidase, cytokinin-activating enzymes).

Secondary pathways were also highly represented and referred to carotenoid, triterpene, and flavonoid pathways (Fig. 3d). For instance, flavonoid synthases or isomerases were more diverse and overexpressed in at least 50% of the plant species. Enzymes involved in diverse reactions covered phenylpropanoid and carotenoid pathways, such as carotene hydroxylases and carotenoid dioxygenases. Furthermore, we observed an overexpression of reactions linked to polyamine biosynthesis and purine degradation processes. Other enzymes such as γ-aminobutyrate aminotransferase, spermidine synthase, and transaminases were overexpressed in 47% of the species (Tab. S5). These results suggested the need for improved nitrogen resource processes and remobilization to thrive in such extreme conditions. In contrast, Atacama plants have likely contracted or negatively regulated gene families involved in energy processes and certain terpene pathways (Fig. S4, Tab. S4 and S5).

### Convergent biochemical strategies in specific vegetation belts

Prepuna species strongly regulated chain length and saturation level of fatty acids as well as cutin and suberin biosynthesis through the overexpression of dedicated enzymes (*e.g.* very long chain reductases and hydroxycinnamoyl transferase). Nitrogen uptake and processing were also highlighted, for instance with the overexpression of glutaminases (Fig. 3d). Conversely, reactions catalyzed by succinate semialdehyde and lariciresinol reductases or by epimerases were overexpressed (and expanded to a lower extent) in Steppe species. Overall, the comparison of the number of expressed gene isoforms and gene expression levels between Atacama and Sister species highlighted tremendous convergences of biochemical strategies. The analysis of these genetic evolutions unveiled relevant management of resources (*i.e.* carbon and nitrogen). Also, several reactions were specifically more diverse and overexpressed in Steppe or Prepuna species to satisfy environmental demands.

### Genetic legacies and positively selected genes are shaped by environmental constraints

To test whether entire biochemical pathways were enriched as compared to related Sister species, we performed Fisher’s exact tests (Wieder *et al.,* 2021 and Fig. 1). Enriched pathways linked to the synthesis of protective compounds were over-represented and greatly influenced by environmental constraints (Fig. S5, Tab. S7 and S8). For instance, we observed a significant enrichment of primary pathways involved in proline, inositol and fatty acyl biosynthesis in Prepuna species. Conversely, biochemical pathways characterized as enriched in Steppe species referred to the synthesis of phenolics, quaternary ammonium and cyanogenic glycoside compounds.

To get insights into the variations in adaptive biochemical strategies employed in different vegetation belts, we conducted a reaction and pathway enrichment analysis comparing the 13 Steppe to the 13 Prepuna species (Fig. 4, S6 and S7, Tab. S9 and S10). Results confirmed (i) the existence of similarities and divergences among the biochemical strategies adopted by plants to respond to major environmental constraints and (ii) a significant homogeneity of enriched reactions and pathways across the different Prepuna or Steppe species. Overall, these findings identified multiple evolutionary convergences illustrated by the high proportion of diverse reactions and overexpressed in more than 50% of the plant species (Fig. 3 and 4).

**Fig. 4.**
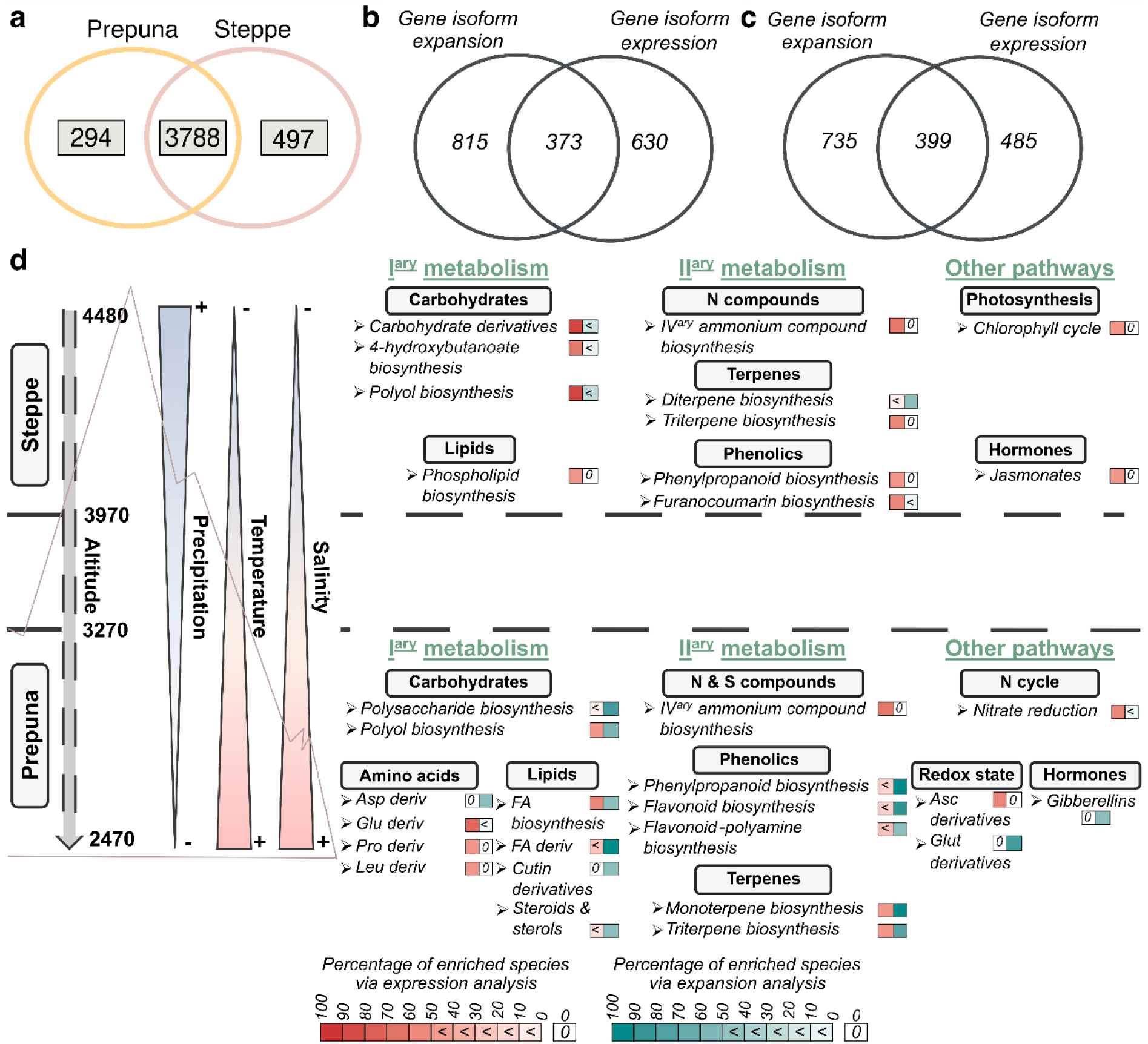
Reaction enrichment analysis comparing Steppe and Prepuna species. **a.** Venn diagram describing the distribution of the predicted annotated reactions in the 13 Steppe species and 13 Prepuna species. **b.** Global trends in the distribution of expanded (*i.e.* diverse) or overexpressed reactions in at least one Steppe species. **c.** Global trends in the distribution of diverse or overexpressed reactions in at least one Prepuna species. **d.** Depiction of the reactions expanded or overexpressed in at least 50% of the species from Steppe or Prepuna (*i.e.* at least 7 species). *Asp*: aspartate, *B-H*: Benjamini-Hochberg, *Chl*: chlorophyll, *Deriv:* derivatives, *Glu*: glutamate, *Leu*: leucine, *Pro*: proline.

Finally, we tested whether enriched reactions were also positively selected in the Atacama species. For this purpose, we tested for the existence of PSGs from distinct ortholog groups within Prepuna or Steppe vegetation belts, while previous analyses focused on PSGs in Atacama *versus* Sisters (Eshel et al., 2021). Significant signs of positive selection to a vegetation belt included 13 ortholog groups for the Prepuna and 23 ortholog groups for the Steppe (Tab. S12). Relevant to their association with stressful environments, GO terms of these ortholog groups included “response to UV and light stimulus” (Prepuna) and “response to stress (*e.g.* light, temperature)” (Steppe). In addition, we found the vegetation belt PSGs included genes assigned the dimethylphylloquinone dehydrogenase (RXN-17007, Prepuna) and a putative anthranilate methyltransferase (RXN-14453, Steppe) reactions, where each reaction was also more likely to be diverse in Atacama species than Sisters (Tab. S11 and S12).

### Correlation between enriched reactions and adaptive metabolites

The enriched reactions also coincide with the potential regulation of a previously discovered set of 39 metabolites employed by plants to cope with the extreme conditions of the Atacama (Dussarrat *et al.,* 2022, Tab. S11). 44% (17 of the 39) of metabolites that better track altitude in the Atacama transect (herein sentinel metabolites, Dussarrat *et al.,* 2022) were represented in the expanded or overexpressed reactions in Atacama plant species (Fig. S8). Hence, the next objective was to relate enriched reactions to sentinel metabolites and explore the link between gene diversity and metabolic dynamics with regard to elevation. We hypothesized that enriched reactions correlated with sentinel metabolites could be involved in adaptation to the highly contrasting environmental conditions observed across the Atacama desert transect. A total of 16 species (nine plant families), were used in both transcriptomics and metabolomics analyses (Fig. 5a). For these species, correlations between sentinel metabolites and reactions enriched in at least 50% of the species were tested. A similar test was performed using a less stringent enrichment threshold (*i.e.* reactions enriched in at least 3 times more Atacama species than Sisters, Tables S13 and S14). A total of 1,493 and 1,002 correlations (Pearson *P* < 0.05 and *r* ≥ 0.7) between the mean normalized intensity of the metabolic predictors and the number of expressed gene isoforms or top-expressed gene isoforms per reaction, respectively. Within these lists, 457 reaction gene copy number correlations and 417 top-expressed reaction number correlations were also phylogenomically enriched in Atacama *versus* Sisters from either phylogenomic or gene expression expansion analysis (Tab. S14). Metabolites that correlated with the most Atacama-enriched reactions were D-alanyl-d-alanine, quercetin 3-(6’’-ferulylglucoside) and cucurbitoside F (104, 85 and 69 correlations, respectively, Fig. 5b). In addition, multiple Atacama-enriched reactions had a related function to their correlated putative metabolites. For instance, the triterpene cucurbitoside F accumulated in the grass *Aristida adscensionis* and the aster *Ambrosia artemisioides* (Tab. S14, Fig. S9a), which are the two species with the highest copy numbers of a reaction in triterpene biosynthesis (RXN-9631, Fig. S9b). Similarly, tri-coumaroyl spermidine was accumulated in *Solanum chilense* and *Fagonia chilense*, which had the greatest number of top-expressed gene isoforms in reactions that participate in coumarate metabolism (RXN-21007, RXN-7575, and RXN-13959).

**Fig. 5.**
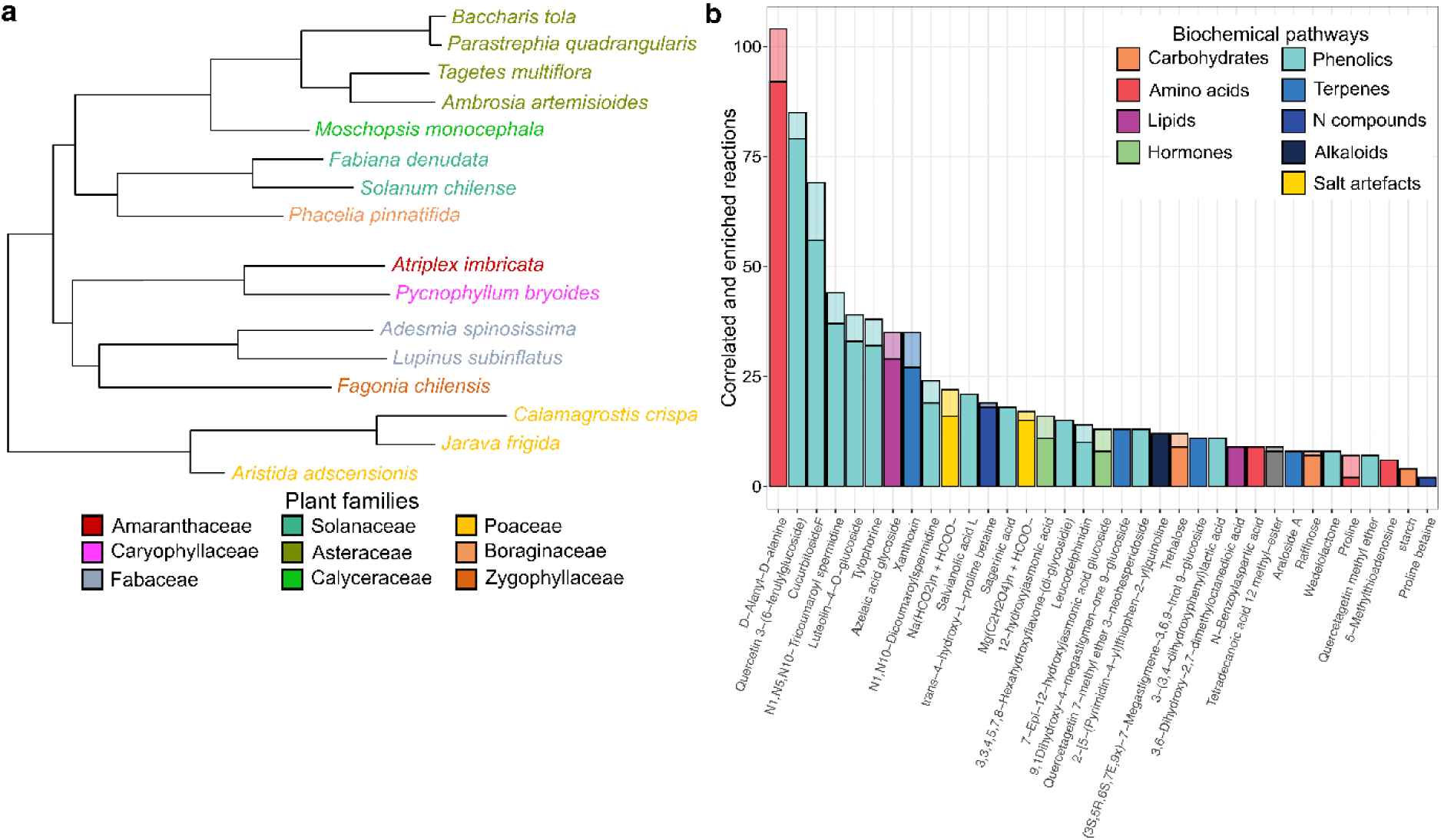
Correlation between enriched reactions and metabolites predicting plant environment in the Atacama Desert. **a.** Sixteen species of plants with both transcriptome annotation and metabolomics. The tree was adapted from Eshel et al., 2021. Species colour refers to taxonomic group. **b.** Number of Atacama-enriched reactions that correlate (Pearson r ≥ 0.7) with the average intensity per species of each metabolic predictor (Dussarrat et al., 2022). Reactions enriched in either gene isoform diversity or expression analyses were included in this graph. Colours refer to the biochemical class of the metabolite, where lighter shades represent the high convergence set (reactions enriched in at least 50% of the species) and darker shades represent a less stringent threshold, where reactions are only enriched in 3 or more Atacama species compared to Sisters (See Tab. S13).

In addition to Pearson’s correlation, we ranked important reactions for predicting metabolite concentrations by machine learning. This procedure could reveal predictive reaction-to-metabolite relationships that are not linearly correlated with metabolite concentration, that we could not find with our previous analysis. In addition, cross-validation of the machine learning lowered the regression score for colinear relationships where reactions occur in only a few species. For instance, high importance was assigned to the reaction 1-Phosphatidylinositol-kinase-Rxn, which generally predicted metabolites that accumulate in *Atriplex imbricata.* Overall, Tab. S14 provides examples of links between shared enriched reactions and predictive metabolites that are likely to allow plants to thrive in the extreme conditions of the Atacama Desert.

## Discussion

### Evolution led to generic metabolic strategies in extreme plants from diverse lineages

How plants adapt to their environment has been of great interest since the domestication of plants in the harsh environments of the Fertile Crescent 10,000 years ago (Riehl et al., 2012). Presumably, the evolution process by natural selection should display some of the most efficient genetic traits for plant survival under harsh conditions. We interrogated the metabolic mechanisms underlying adaptive strategies of multiple Atacama plant species. We compared gene expansion and expression levels at the genome-scale between 32 Atacama species and 32 related Sisters, discovering convergent and divergent biochemical pathways relevant to plant survival. Such a comparative approach allowed us to investigate the plant response to major abiotic constraints while preserving the ecological and evolutionary context (Kang et al., 2020). Results showed convergent evolutions of pre-existing reactions and pathways in response to the main environmental pressures encountered in the Atacama Desert.

First, Atacama and Sister species were not so fundamentally distinct and the extreme majority (99%) of the 463 exclusive reactions in Atacama plants were observed in only one, two or three species. Atacama plants also conserved most of the biochemical reservoir from their ancestors since only 6% of the total annotated reactions were unique to Sisters. While phylogenomics and metabolomics approaches permit the detection of these unknown traits, the majority of genes and metabolites highlighted as relevant for the adaption of Atacama species pertained to conserved processes (Eshel et al., 2021; Dussarrat et al., 2022). Considering the occurrence of vital genome innovations over the past 145 million years of angiosperm evolution, twelve million years (*i.e*. age of the Atacama Desert) represents a short time-scale for the development of new species-specific reactions allowing survival of 14 distinct plant lineages (Jordan et al., 2014; Benton et al., 2021). Nevertheless, these exclusive reactions might have a role in improving plant fitness at the species level, which could be evaluated through an untargeted comparison of adapted and non-adapted species from the same genus.

Conversely, a significant number of biochemical convergences was shown as a result of the plant evolution process in the Atacama Desert among conserved reactions between Atacama and Sister species. These shared mechanisms demonstrated a high potential to modulate resource uptake and allocation between development and stress tolerance. Modulation of major metabolic processes was targeted by expanded and overexpressed reactions in at least 50% of the species studied. Genes underlying these enriched biochemical routes also pointed to the regulation of metabolites employed by Atacama plants to face extreme environmental conditions (Dussarrat et al., 2022), such as proline, jasmonic acids, quercetin, and tricoumaroyl spermidine compounds as well as carotenoid cleavage and chlorophyll cycle. With carotenoid and chlorophyll-related pathways, quercetin was the most frequently observed compound in exclusive, expanded, and overexpressed reactions. This flavonoid was linked to various roles as an antioxidant, a mediator of interaction with nitrogen-fixing bacteria, and its links with both hormones (*e.g.* abscisic acid) and redox buffers (*e.g.* glutathione) (Singh et al., 2021). The significance of these links between enriched reactions and adaptive metabolites was further tested, yielding a consequent number of correlations. Both Pearson’s correlation and machine learning pointed to relevant reaction-to-metabolite relationships such as links between the 1-Phosphatidylinositol-kinase-Rxn and metabolites from *Atriplex imbricata,* a genus known to accumulate salt in its leaves (Belkheiri and Mulas, 2013). Overall, our findings reinforce the place of gene duplication as an evolutionary mechanism to accumulate adaptive metabolites (Flagel and Wendel, 2009; Méteignier et al., 2022).

Hence, while adaptation was thought of as mainly species-dependent (Turner, 2018; Dussarrat et al., 2021), these findings strongly suggest that a relevant part of this adaptation is conversely the result of convergent evolutions of regulatory processes. In other words, the development of adaptive traits providing a selective advantage is more likely to occur from the regulation of pre-existing compounds and pathways than from the emergence of new ones. Furthermore, most of these convergences represented conserved compounds among wild and crop species. Finally, two independent analytical approaches highlighted many of these metabolic traits (Dussarrat et al., 2022), which thus hold great promise for potential crop engineering.

### Intense environmental pressures define the evolutionary trajectories

Solar irradiance, water and nitrogen scarcity are three critical parameters for plant survival in the Atacama Desert (Eshel et al., 2021). The Steppe undergoes negative daily temperatures while Prepuna species are subjected to extreme salinity (Díaz et al., 2019). Our study unveiled several adaptive mechanisms that have been fixed through evolution and answered a significant proportion of these challenging environmental constraints.

Carbon is not a limiting resource in desertic regions due to intense solar irradiance (Eshel et al., 2021). Enriched reactions in Atacama plants were linked to the regulation of chlorophyll level and chlorophyll a/chlorophyll b ratio, suggesting the regulation of energy capture to mitigate oxidative stress, in agreement with other extreme plants (Sato et al., 2015; Cui et al., 2019). The allocation of carbon reserves between plant growth and defense seemed finely tuned through evolution with, for example, the modulation of gibberellin and cytokinin pathways (Fig 3d). In addition, enriched reactions in at least 50% were not limited to secondary pathways, reinforcing the role of primary metabolism in plant resilience to harsh climates (Dussarrat et al., 2022). Synthesis of oligosaccharides and polyols could be linked to low water availability (Williamson et al., 2002; Pamuru et al., 2021). Reactions and pathways involved in lipid metabolism were also universally represented. Synthesis of waxes and long fatty acyl chains were previously highlighted in the Atacama plants and can refer to how plants limit evapotranspiration and minimize the impact of solar irradiance or osmotic stress (Kolattukudy, 1970; Frugone-Álvarez et al., 2023). Furthermore, the extensive expansion and overexpression of reactions involved in the control of chain length and saturation level of fatty acyls could support the role of membrane fluidity in adaptation (Li et al., 2020). Finally, these changes accompanied the regulation of the TCA cycle and proteinogenic amino acid pathways, two major precursors of secondary compounds (Yang et al., 2020). Thus, a significant part of the fixed carbon was likely to be used to produce secondary protective compounds. The presence of the mevalonate pathway and carotenoid synthesis and cleavage processes among the reactions expanded and overexpressed in more than 50% of the Atacama species is consistent with their role in stress mitigation and their links with hormonal and redox pathways (Havaux, 2014). Nitrogen-related compounds (*e.g.* polyamines, quaternary ammonium compounds) and phenolics are other metabolic features employed by plants to face a myriad of climate constraints (Kinnersley and Turano, 2000; Martinelli et al., 2007; Dussarrat et al., 2021). A significant proportion of the enriched reactions is directly or indirectly linked to redox homeostasis through reactive oxygen species scavenging (*e.g.* polyphenols) or proline and carotenoid cleavage respectively, supporting its central place in adaptation.

Atacama plant transcriptomes strongly reflect the specific requirements angered by the elevation (Fig. 3d, 4 and 5). This was exemplified by the over-representation of enriched reactions and pathways linked to polyol and sterol synthesis in Prepuna, where drought and salinity are extremely severe (Rogowska and Szakiel, 2020; Eshel et al., 2021). Nitrogen starvation is of major importance in this vegetation belt, illustrated by the extensive genetic expansion and expression of reactions linked to plant defense mechanisms through the monitoring of glutamine, glutamate, GABA and polyamine pathways (Solomon and Oliver, 2002; Martinelli et al., 2007). In addition, evolution led to an increased synthesis of some flavonoid-polyamine conjugates (*e.g.* tricoumaroyl spermidine) whose roles remain poorly described (Dussarrat et al., 2021). In contrast, Steppe species favoured the production of a plethora of secondary compounds (Fig. 3d and 5). Carotenoids and other isoprenoids could be used for their advantages to mitigate abiotic constraints (Havaux, 2014). Steppe species were likely to favour the tolerance to freezing stress via the synthesis of lignans and the tolerance to oxidative stress through 4-hydroxybutanoate accumulation and pyridine nucleotide repair (via epimerases), which were recently considered as a relevant player in NAD(P)H metabolism (Breitkreuz et al., 2003; Gakière et al., 2018). Plants from higher levels also tend to enrich reactions involved in producing a wide range of metabolites engaged in biotic defense such as coumarins and cyanogenic glucosides (Gleadow and Woodrow, 2002; Stringlis et al., 2019).

Although great successes in plant breeding, yields of the best ideotypes are threatened by climate change (Seneviratne et al., 2012; Bailey-Serres et al., 2019). We thus need innovative research strategies that depart from the current reductionist single-species approach to discover universal plant resilience mechanisms, which should be more easily transferable to crops. This transcriptome-scale comparative analysis provided a unique goldmine of genetic targets for engineering resilient crops against major abiotic threats. Interrogating Atacama plant evolution has shown that extreme plant uniqueness lies in the regulation of pre-existing pathways and has unveiled a high degree of convergences between biochemical strategies selected to face harsh climate conditions. These generic strategies were relevant to plant resilience against osmotic (*e.g.* drought and salinity), frost and high irradiance stress as well as low soil suitability. Hence, these findings pave the way for wider use of these generic metabolic mechanisms to provide sustainable solutions to improve global food security. A thrilling perspective will be to investigate whether these generic biochemical reactions are also involved in adaptation to other extreme lands.

## Author contributions

RG and CL conceived the Atacama project. TD, RNP, FPD, GMC, YG, DR, CL, PP and RG participated in the conceptualization of this enrichment analysis. TD and RNP produced the initial annotated reactions and pathways table. TD, RNP, TCM and SP performed the enrichment analyses. TLJ performed the correlation analysis between enriched reactions and metabolic intensities. TLJ, LA, VMS, DS, and BS explored positively selected genes. Biological interpretation of the results was conducted by TD, RNP, TCM, SP, FPD, VA, CL, PP and RG. TD, RNP, PP and RG wrote the manuscript with feedback from all co-authors.

## Acknowledgements

We acknowledge the Comunidad de Talabre for providing us access to the Talabre-Lejía transect. We also thank Gil Eshel and Fabien Jourdan for their help and analytical advice. We thank Fondo Nacional de Desarrollo Científico y Tecnológico (FONDECYT 1180759), DOI EVONET Proyect and ANID – Millennium Science Initiative Program – iBio ICN17_022 as well as Fondo de Desarrollo de Áreas Prioritarias (FONDAP) Center for Genome Regulation (15200002) for financial support to R. A. Gutiérrez. We acknowledge the US Department of Energy Biological and Environmental Research (DOE-BER) Grant DE SC0014377 to G. M. Coruzzi and R. A. Gutiérrez, and the Zegar Family Foundation (A16-0051) to G. M. Coruzzi. We thank Fondecyt Iniciación 11150107 for financial support to R. Nilo-Poyanco. F. P. Díaz and C. Latorre thank ANID - Millennium Science Initiative Program Nucleus AFOREST (NCS2022_24) and IEB (ANID FB210006). P. Pétriacq and Y. Gibon thank MetaboHUB (ANR-11-INBS-0010) for financial support.

## Data availability

All data and metadata are available in supplemental tables. In addition, transcriptomic data (*i.e.* top 10% expressed genes from Atacama and related species) are available in Eshel et al., 2021 and metabolomics data (intensity of the metabolic features) are available in Dussarrat et al., 2022.

## Abbreviations

ORA: over-representation analysis
PSGs: positively selected genes

## References

Báez S, Collins SL (2008) Shrub invasion decreases diversity and alters community stability in Northern Chihuahuan Desert plant communities. PLoS ONE 3: e2332

Bailey-Serres J, Parker JE, Ainsworth EA, Oldroyd GED, Schroeder JI (2019) Genetic strategies for improving crop yields. Nature 575: 109–118

Belkheiri O, Mulas M (2013) The effects of salt stress on growth, water relations and ion accumulation in two halophyte *Atriplex* species. Environmental and Experimental Botany 86: 17–28

Benjamini Y, Hochberg Y (1995) Controlling the false discovery rate: a practical and powerful approach to multiple testing. Journal of the Royal Statistical Society: Series B (Methodological) 57: 289–300

Benton MJ, Wilf P, Sauquet H (2021) The Angiosperm terrestrial revolution and the origins of modern biodiversity. New Phytologist 233: 2017–2035

Bolger A, Scossa F, Bolger ME, Lanz C, Maumus F, Tohge T, Quesneville H, Alseekh S, Sørensen I, Lichtenstein G, et al (2014) The genome of the stress-tolerant wild tomato species *Solanum pennellii*. Nature Genetics 46: 1034–1038

Breitkreuz KE, Allan WL, Van Cauwenberghe OR, Jakobs C, Talibi D, André B, Shelp BJ (2003) A novel γ-hydroxybutyrate dehydrogenase. Journal of Biological Chemistry 278: 41552–41556

Carrasco-Puga G, Díaz FP, Soto DC, Hernández-Castro C, Contreras-López O, Maldonado A, Latorre C, Gutiérrez RA (2021) Revealing hidden plant diversity in arid environments. Ecography 44: 98–111

Caspi R, Billington R, Keseler IM, Kothari A, Krummenacker M, Midford PE, Ong WK, Paley S, Subhraveti P, Karp PD (2020) The MetaCyc database of metabolic pathways and enzymes - a 2019 update. Nucleic Acids Research 48: D445–D453

Chae L, Kim T, Nilo-Poyanco R, Rhee SY (2014) Genomic signatures of specialized metabolism in plants. Science 344: 510–513

Chen T, Guestrin C (2016) XGBoost: A scalable tree boosting system. Proceedings of the 22nd ACM SIGKDD international conference on knowledge discovery and data mining. ACM, San Francisco California USA, pp 785–794

Cui G, Ji G, Liu S, Li B, Lian L, He W, Zhang P (2019) Physiological adaptations of *Elymus dahuricus* to high altitude on the Qinghai–Tibetan plateau. Acta Physiologiae Plantarum 41: 115

Defossez E, Pitteloud C, Descombes P, Glauser G, Allard P-M, Walker TWN, Fernandez-Conradi P, Wolfender J-L, Pellissier L, Rasmann S (2021) Spatial and evolutionary predictability of phytochemical diversity. Proceedings of the National Academy of Sciences USA 118: e2013344118

Díaz FP, Frugone M, Gutiérrez RA, Latorre C (2016) Nitrogen cycling in an extreme hyperarid environment inferred from δ^15^N analyses of plants, soils and herbivore diet. Scientific Reports 6: 22226

Díaz FP, Latorre C, Carrasco-Puga G, Wood JR, Wilmshurst JM, Soto DC, Cole TL, Gutiérrez RA (2019) Multiscale climate change impacts on plant diversity in the Atacama Desert. Global Change Biology 25: 1733–1745

Dussarrat T, Decros G, Díaz FP, Gibon Y, Latorre C, Rolin D, Gutiérrez RA, Pétriacq P (2021) Another tale from the harsh world: how plants adapt to extreme environments. Annual Plant Reviews online. Annual Plant Reviews online 4: pp 551–603

Dussarrat T, Prigent S, Latorre C, Bernillon S, Flandin A, Díaz FP, Cassan C, Van Delft P, Jacob D, Varala K, et al (2022) Predictive metabolomics of multiple Atacama plant species unveils a core set of generic metabolites for extreme climate resilience. New Phytologist nph.18095 234, 1614–1628

Eshel G, Araus V, Undurraga S, Soto DC, Moraga C, Montecinos A, Moyano T, Maldonado J, Díaz FP, Varala K, et al (2021) Plant ecological genomics at the limits of life in the Atacama Desert. Proceedings of the National Academy of Sciences USA 118: e2101177118

Fiehn O (2002) Metabolomics — the link between genotypes and phenotypes. *In* C Town, ed, Functional Genomics. Springer Netherlands, Dordrecht, pp 155–171

Flagel LE, Wendel JF (2009) Gene duplication and evolutionary novelty in plants. New Phytologist 183: 557–564

Frugone-Álvarez M, Contreras S, Meseguer-Ruiz O, Tejos E, Delgado-Huertas A, Valero-Garcés B, Díaz FP, Briceño M, Bustos-Morales M, Latorre C (2023) Hydroclimate variations over the last 17,000 years as estimated by leaf waxes in rodent middens from the south-central Atacama Desert, Chile. Quaternary Science Reviews 311: 108084

Gakière B, Hao J, Bont L de, Pétriacq P, Nunes-Nesi A, Fernie AR (2018) NAD+ biosynthesis and signaling in plants. Critical Reviews in Plant Sciences 37: 259–307

Gleadow RM, Woodrow IE (2002) Mini-review: constraints on effectiveness of cyanogenic glycosides in herbivore defense. Journal of Chemical Ecology 28: 1301–1313

Havaux M (2014) Carotenoid oxidation products as stress signals in plants. Plant Journal 79: 597–606

Jia Q, Kong D, Li Q, Sun S, Song J, Zhu Y, Liang K, Ke Q, Lin W, Huang J (2019) The function of inositol phosphatases in plant tolerance to abiotic stress. IJMS 20: 3999

Jordan TE, Kirk-Lawlor NE, Blanco NP, Rech JA, Cosentino NJ (2014) Landscape modification in response to repeated onset of hyperarid paleoclimate states since 14 Ma, Atacama Desert, Chile. Geological Society of America Bulletin 126: 1016–1046

Joswig JS, Wirth C, Schuman MC, Kattge J, Reu B, Wright IJ, Sippel SD, Rüger N, Richter R, Schaepman ME, et al (2021) Climatic and soil factors explain the two-dimensional spectrum of global plant trait variation. Nature Ecology and Evolution. doi: 10.1038/s41559-021-01616-8

Kanehisa M, Goto S, Sato Y, Kawashima M, Furumichi M, Tanabe M (2014) Data, information, knowledge and principle: back to metabolism in KEGG. Nucleic Acids Research 42: D199–D205

Kang S-H, Pandey RP, Lee C-M, Sim J-S, Jeong J-T, Choi B-S, Jung M, Ginzburg D, Zhao K, Won SY, et al (2020) Genome-enabled discovery of anthraquinone biosynthesis in *Senna tora*. Nature Communications 11: 5875

Karp PD, Midford PE, Billington R, Kothari A, Krummenacker M, Latendresse M, Ong WK, Subhraveti P, Caspi R, Fulcher C, et al (2021) Pathway Tools version 23.0 update: software for pathway/genome informatics and systems biology. Briefings in Bioinformatics 22: 109–126

Kinnersley AM, Turano FJ (2000) Gamma aminobutyric acid (GABA) and plant responses to stress. Critical Reviews in Plant Sciences 19: 479–509

Kolattukudy PE (1970) Plant waxes. Lipids 5: 259–275

Kumari M, Joshi R, Kumar R (2020) Metabolic signatures provide novel insights to *Picrorhiza kurroa* adaptation along the altitude in Himalayan region. Metabolomics 16: 77

Li J, Liu L-N, Meng Q, Fan H, Sui N (2020) The roles of chloroplast membrane lipids in abiotic stress responses. Plant Signaling & Behavior 15: 1807152

Lou Y, Gou J-Y, Xue H-W (2007) PIP5K9, an *Arabidopsis* phosphatidylinositol monophosphate kinase, interacts with a cytosolic invertase to negatively regulate sugar-mediated root growth. The Plant Cell 19: 163–181

Lugan R, Niogret M-F, Leport L, Guégan J-P, Larher FR, Savouré A, Kopka J, Bouchereau A (2010) Metabolome and water homeostasis analysis of *Thellungiella salsuginea* suggests that dehydration tolerance is a key response to osmotic stress in this halophyte. Plant Journal 64: 215–229

Martinelli T, Whittaker A, Bochicchio A, Vazzana C, Suzuki A, Masclaux-Daubresse C (2007) Amino acid pattern and glutamate metabolism during dehydration stress in the “resurrection” plant *Sporobolus stapfianus*: a comparison between desiccation-sensitive and desiccation-tolerant leaves. Journal of Experimental Botany 58: 3037–3046

Méteignier L-V, Nützmann H-W, Papon N, Osbourn A, Courdavault V (2022) Emerging mechanistic insights into the regulation of specialized metabolism in plants. Nature Plants 9: 22–30

Murrell B, Weaver S, Smith MD, Wertheim JO, Murrell S, Aylward A, Eren K, Pollner T, Martin DP, Smith DM, et al (2015) Gene-wide identification of episodic selection. Molecular Biology and Evolution 32: 1365–1371

Pamuru RR, Puli COR, Pandita D, Wani SH (2021) Sugar alcohols and osmotic stress adaptation in plants. *In* SH Wani, MP Gangola, BR Ramadoss, eds, Compatible Solutes Engineering for Crop Plants Facing Climate Change. Springer International Publishing, Cham, pp 189–203

Paradis E, Schliep K (2019) ape 5.0: an environment for modern phylogenetics and evolutionary analyses in R. Bioinformatics 35: 526–528

Pond SLK, Frost SDW, Muse SV (2005) HyPhy: hypothesis testing using phylogenies. Bioinformatics 21: 676–679

R Core Team (2021) R: a language and environment for statistical computing. R Foundation for Statistical Computing, Vienna, Austria

Riehl S, Benz M, Conard NJ, Darabi H, Deckers K, Nashli HF, Zeidi-Kulehparcheh M (2012) Plant use in three pre-pottery Neolithic sites of the northern and eastern Fertile Crescent: a preliminary report. Vegetation History Archaeobotany 21: 95–106

Rogowska A, Szakiel A (2020) The role of sterols in plant response to abiotic stress. Phytochemistry Reviews 19: 1525–1538

Sato R, Ito H, Tanaka A (2015) Chlorophyll b degradation by chlorophyll b reductase under high-light conditions. Photosynthesis Research 126: 249–259

Schläpfer P, Zhang P, Wang C, Kim T, Banf M, Chae L, Dreher K, Chavali AK, Nilo-Poyanco R, Bernard T, et al (2017) Genome-wide prediction of metabolic enzymes, pathways, and gene clusters in plants. Plant Physiology 173: 2041–2059

Seneviratne SI, Nicholls N, Easterling D, Goodess CM, Kanae S, Kossin J, Luo Y, Marengo J, McInnes K, Rahimi M, et al (2012) Changes in climate extremes and their impacts on the natural physical environment. *In* CB Field, V Barros, TF Stocker, Q Dahe, eds, Managing the Risks of Extreme Events and Disasters to Advance Climate Change Adaptation. Cambridge University Press, Cambridge, pp 109–230

Singh P, Arif Y, Bajguz A, Hayat S (2021) The role of quercetin in plants. Plant Physiology and Biochemistry 166: 10–19

Smith MD, Wertheim JO, Weaver S, Murrell B, Scheffler K, Kosakovsky Pond SL (2015) Less is more: an adaptive branch-site random effects model for efficient detection of episodic diversifying selection. Molecular Biology and Evolution 32: 1342–1353

Solomon PS, Oliver RP (2002) Evidence that γ-aminobutyric acid is a major nitrogen source during Cladosporium fulvum infection of tomato. Planta 214: 414–420

Stringlis IA, de Jonge R, Pieterse CMJ (2019) The age of coumarins in plant–microbe interactions. Plant and Cell Physiology 60: 1405–1419

Turner NC (2018) Turgor maintenance by osmotic adjustment: 40 years of progress. Journal of Experimental Botany 69: 3223–3233

Voss-Fels KP, Stahl A, Wittkop B, Lichthardt C, Nagler S, Rose T, Chen T-W, Zetzsche H, Seddig S, Majid Baig M, et al (2019) Breeding improves wheat productivity under contrasting agrochemical input levels. Nature Plants 5: 706–714

Wieder C, Frainay C, Poupin N, Rodríguez-Mier P, Vinson F, Cooke J, Lai RP, Bundy JG, Jourdan F, Ebbels T (2021) Pathway analysis in metabolomics: pitfalls and best practice for the use of over-representation analysis. PLoS Comput Biol. 2021 Sep 7;17(9):e1009105.

Williamson JD, Jennings DB, Guo W-W, Pharr DM, Ehrenshaft M (2002) Sugar alcohols, salt stress, and fungal resistance: polyols—multifunctional plant protection? Journal of the American Society for Horticultural Science 127: 467–473

Wishart DS, Feunang YD, Marcu A, Guo AC, Liang K, Vázquez-Fresno R, Sajed T, Johnson D, Li C, Karu N, et al (2018) HMDB 4.0: the human metabolome database for 2018. Nucleic Acids Research 46: D608–D617

Yang T, Li H, Tai Y, Dong C, Cheng X, Xia E, Chen Z, Li F, Wan X, Zhang Z (2020) Transcriptional regulation of amino acid metabolism in response to nitrogen deficiency and nitrogen forms in tea plant root (Camellia sinensis L.). Scientific Reports 10: 6868

Yano R, Nakamura M, Yoneyama T, Nishida I (2005) Starch-related *α* -glucan/water dikinase is involved in the cold-induced development of freezing tolerance in Arabidopsis. Plant Physiology 138: 837–846

Yu G (2022) Data integration, manipulation and visualization of phylogenetic trees, 1st ed. doi: 10.1201/9781003279242

Zhang X, Gu S, Zhao X, Cui X, Zhao L, Xu S, Du M, Jiang S, Gao Y, Ma C, et al (2010) Radiation partitioning and its relation to environmental factors above a meadow ecosystem on the Qinghai-Tibetan Plateau. Journal of Geophysical Research 115: D10106

Ziaco E, Truettner C, Biondi F, Bullock S (2018) Moisture-driven xylogenesis in *Pinus ponderosa* from a Mojave Desert mountain reveals high phenological plasticity: moisture-driven xylogenesis in *Pinus ponderosa*. Plant Cell Environment 41: 823–836

