## Supplemental Figure 1 for "Phylogenetically diverse wild plant species use common biochemical strategies to thrive in the Atacama Desert"

### Slide 1
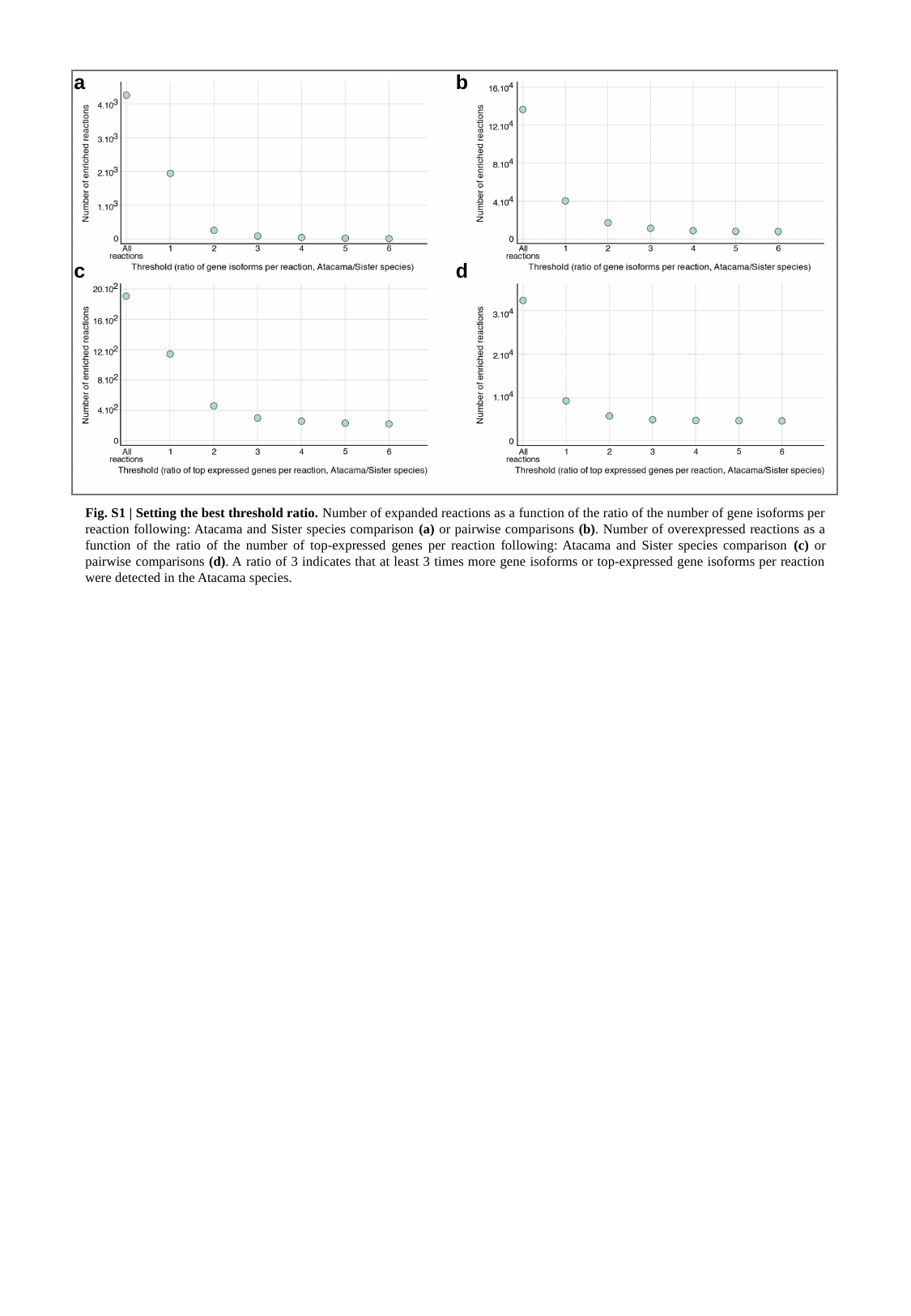

a
b
c
d
Fig. S1 | Setting the best threshold ratio. Number of expanded reactions as a function of the ratio of the number of gene isoforms per reaction following: Atacama and Sister species comparison (a) or pairwise comparisons (b). Number of overexpressed reactions as a function of the ratio of the number of top-expressed genes per reaction following: Atacama and Sister species comparison (c) or pairwise comparisons (d). A ratio of 3 indicates that at least 3 times more gene isoforms or top-expressed gene isoforms per reaction were detected in the Atacama species.
