## Supplemental Figure 2 for "Phylogenetically diverse wild plant species use common biochemical strategies to thrive in the Atacama Desert"

### Slide 1
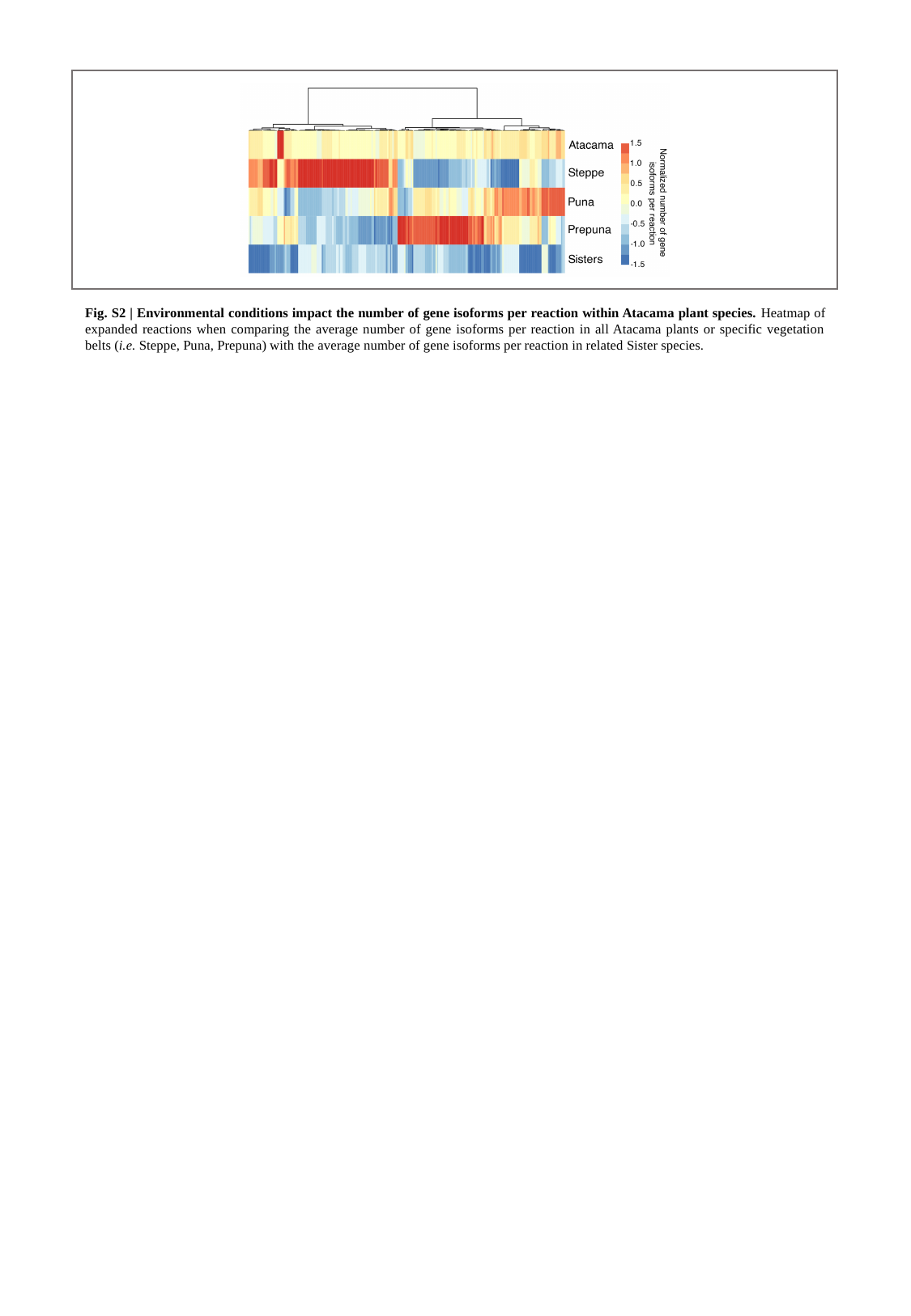

Fig. S2 | Environmental conditions impact the number of gene isoforms per reaction within Atacama plant species. Heatmap of expanded reactions when comparing the average number of gene isoforms per reaction in all Atacama plants or specific vegetation belts (i.e. Steppe, Puna, Prepuna) with the average number of gene isoforms per reaction in related Sister species.
