## Supplemental Figure 3 for "Phylogenetically diverse wild plant species use common biochemical strategies to thrive in the Atacama Desert"

### Slide 1
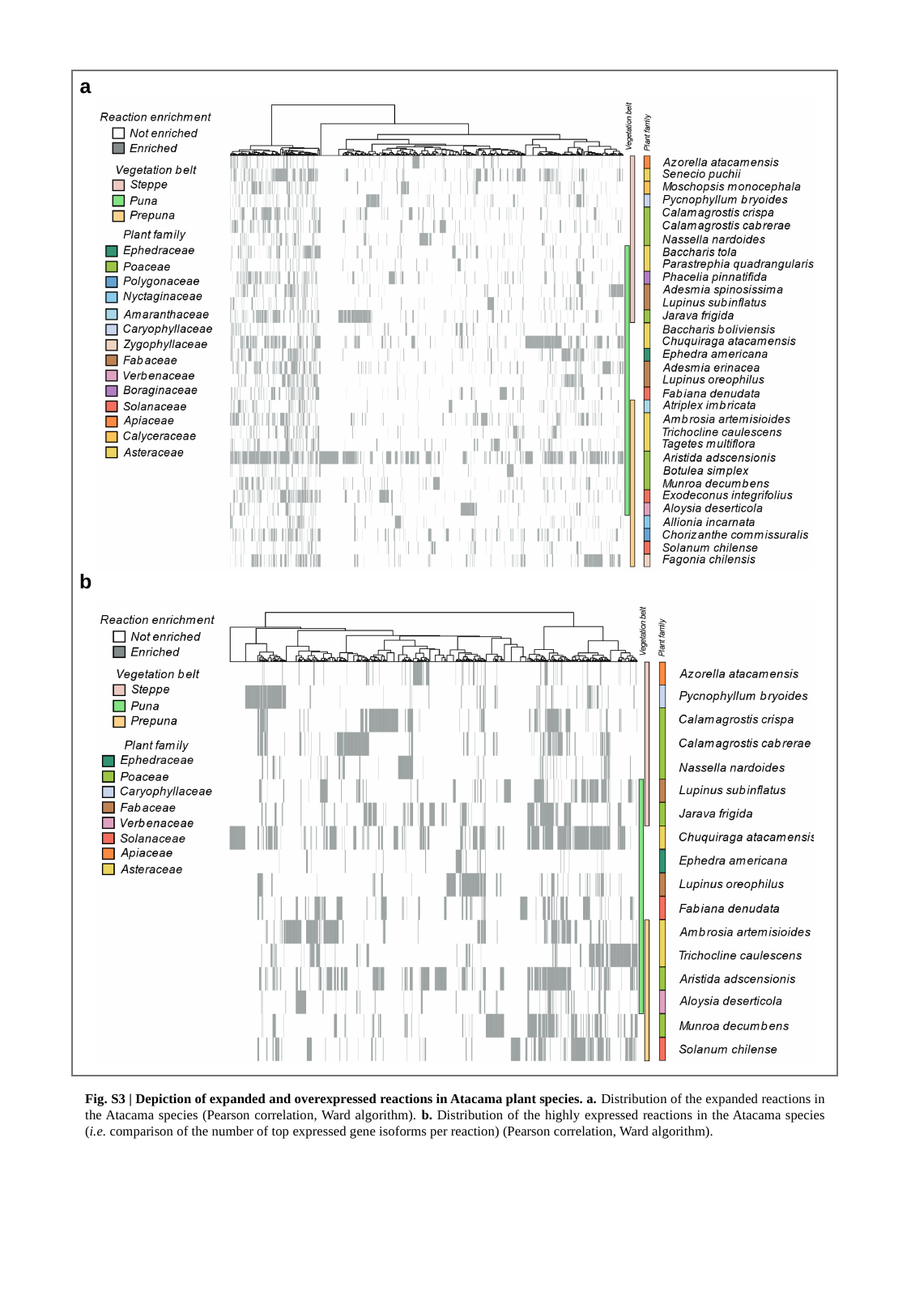

a
b
Fig. S3 | Depiction of expanded and overexpressed reactions in Atacama plant species. a. Distribution of the expanded reactions in the Atacama species (Pearson correlation, Ward algorithm). b. Distribution of the highly expressed reactions in the Atacama species (i.e. comparison of the number of top expressed gene isoforms per reaction) (Pearson correlation, Ward algorithm).
