## Supplemental Figure 4 for "Phylogenetically diverse wild plant species use common biochemical strategies to thrive in the Atacama Desert"

### Slide 1
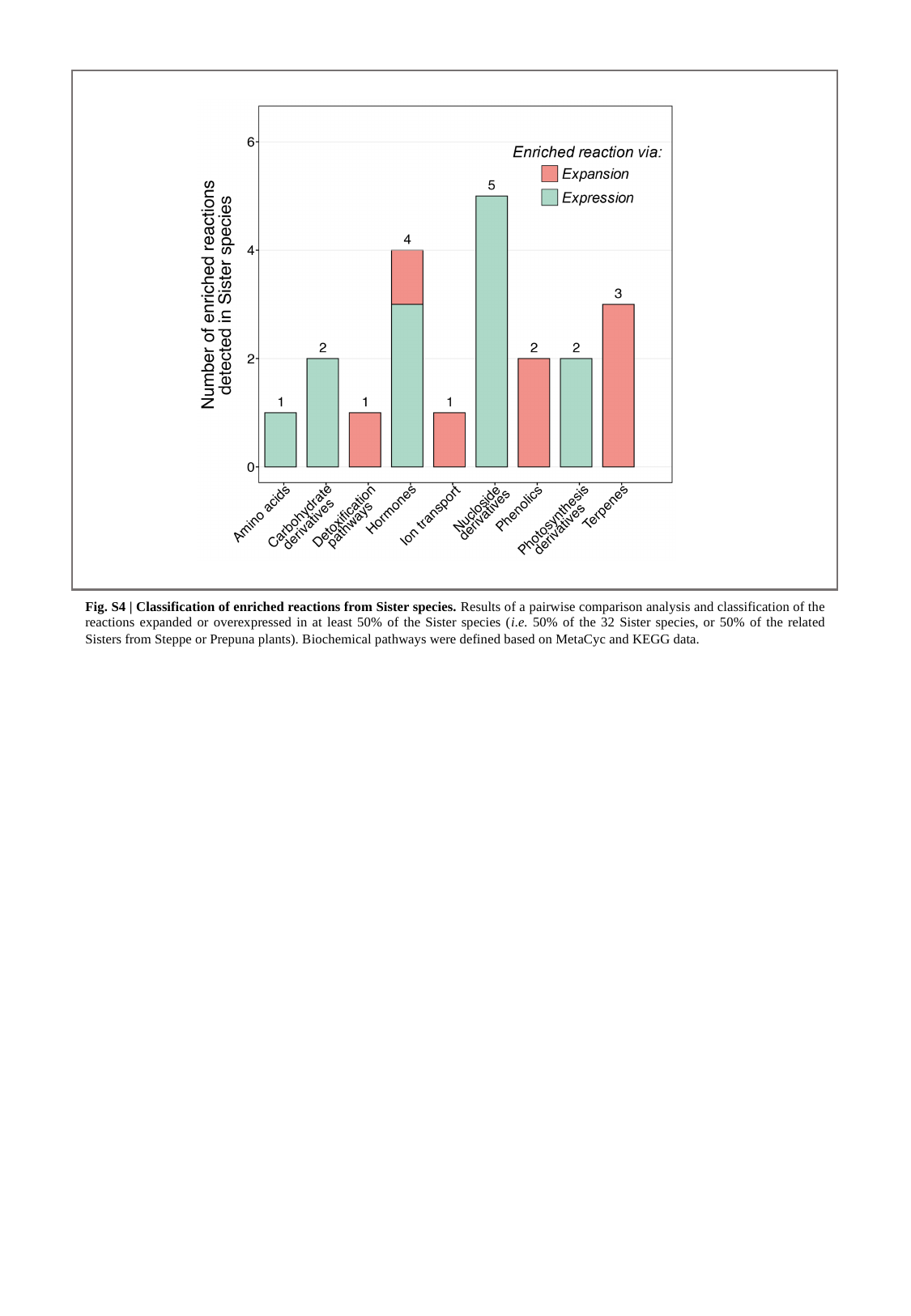

Fig. S4 | Classification of enriched reactions from Sister species. Results of a pairwise comparison analysis and classification of the reactions expanded or overexpressed in at least 50% of the Sister species (i.e. 50% of the 32 Sister species, or 50% of the related Sisters from Steppe or Prepuna plants). Biochemical pathways were defined based on MetaCyc and KEGG data.
