## Supplemental Figure 5 for "Phylogenetically diverse wild plant species use common biochemical strategies to thrive in the Atacama Desert"

### Slide 1
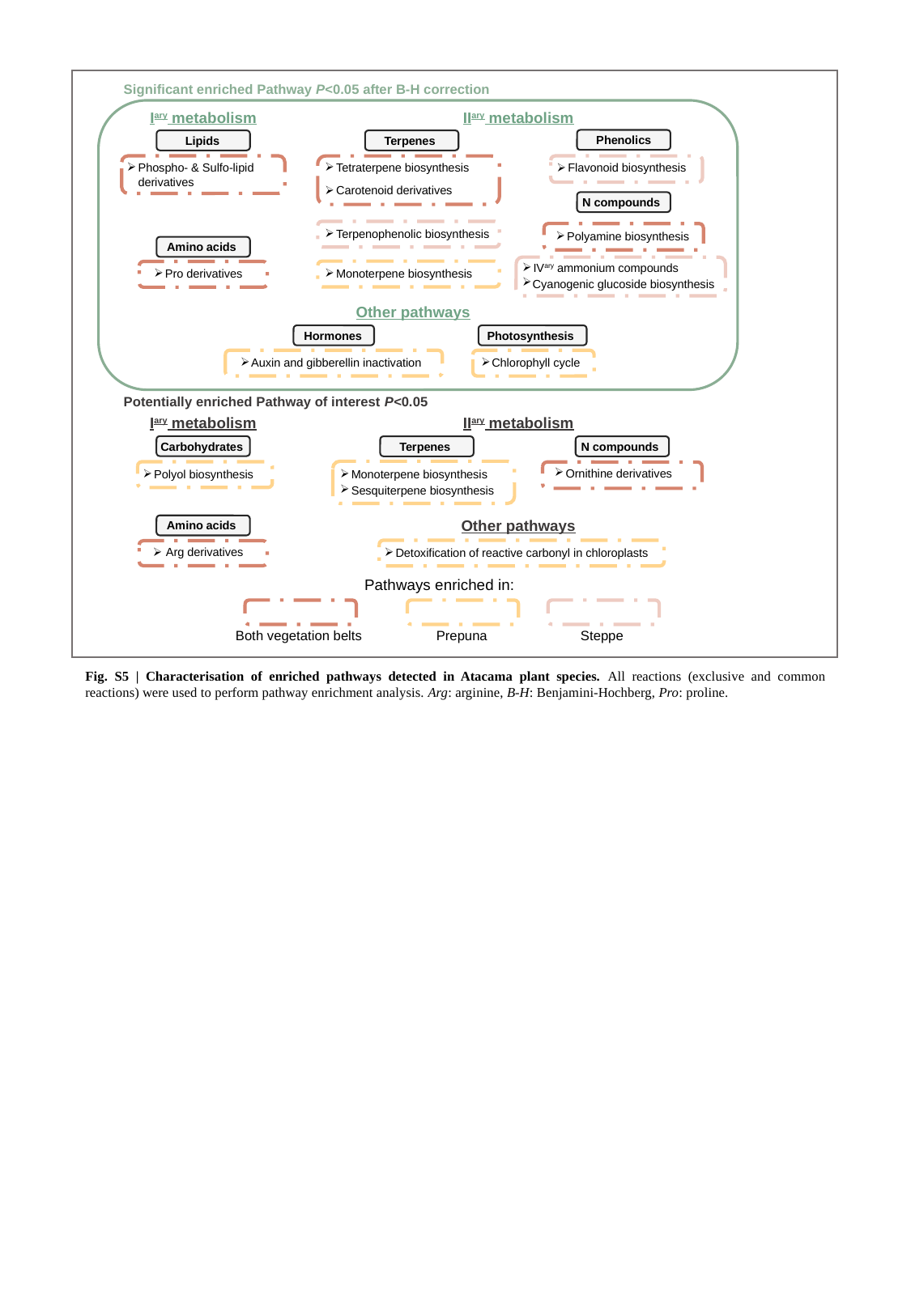

Significant enriched Pathway P<0.05 after B-H correction
I^ary^ metabolism
II^ary^ metabolism
Phenolics
Lipids
Terpenes
 Phospho- & Sulfo-lipid
 derivatives
 Tetraterpene biosynthesis
 Carotenoid derivatives
 Flavonoid biosynthesis
N compounds
 Terpenophenolic biosynthesis
 Polyamine biosynthesis
Amino acids
 IV^ary^ ammonium compounds
 Cyanogenic glucoside biosynthesis
 Monoterpene biosynthesis
 Pro derivatives
Other pathways
Hormones
 Auxin and gibberellin inactivation
Photosynthesis
 Chlorophyll cycle
Potentially enriched Pathway of interest P<0.05
I^ary^ metabolism
II^ary^ metabolism
Carbohydrates
Terpenes
 Monoterpene biosynthesis
 Sesquiterpene biosynthesis
N compounds
 Ornithine derivatives
 Polyol biosynthesis
Other pathways
Amino acids
 Arg derivatives
 Detoxification of reactive carbonyl in chloroplasts
Pathways enriched in:
Both vegetation belts
Prepuna
Steppe
Fig. S5 | Characterisation of enriched pathways detected in Atacama plant species. All reactions (exclusive and common reactions) were used to perform pathway enrichment analysis. Arg: arginine, B-H: Benjamini-Hochberg, Pro: proline.
