## Supplemental Figure 6 for "Phylogenetically diverse wild plant species use common biochemical strategies to thrive in the Atacama Desert"

### Slide 1
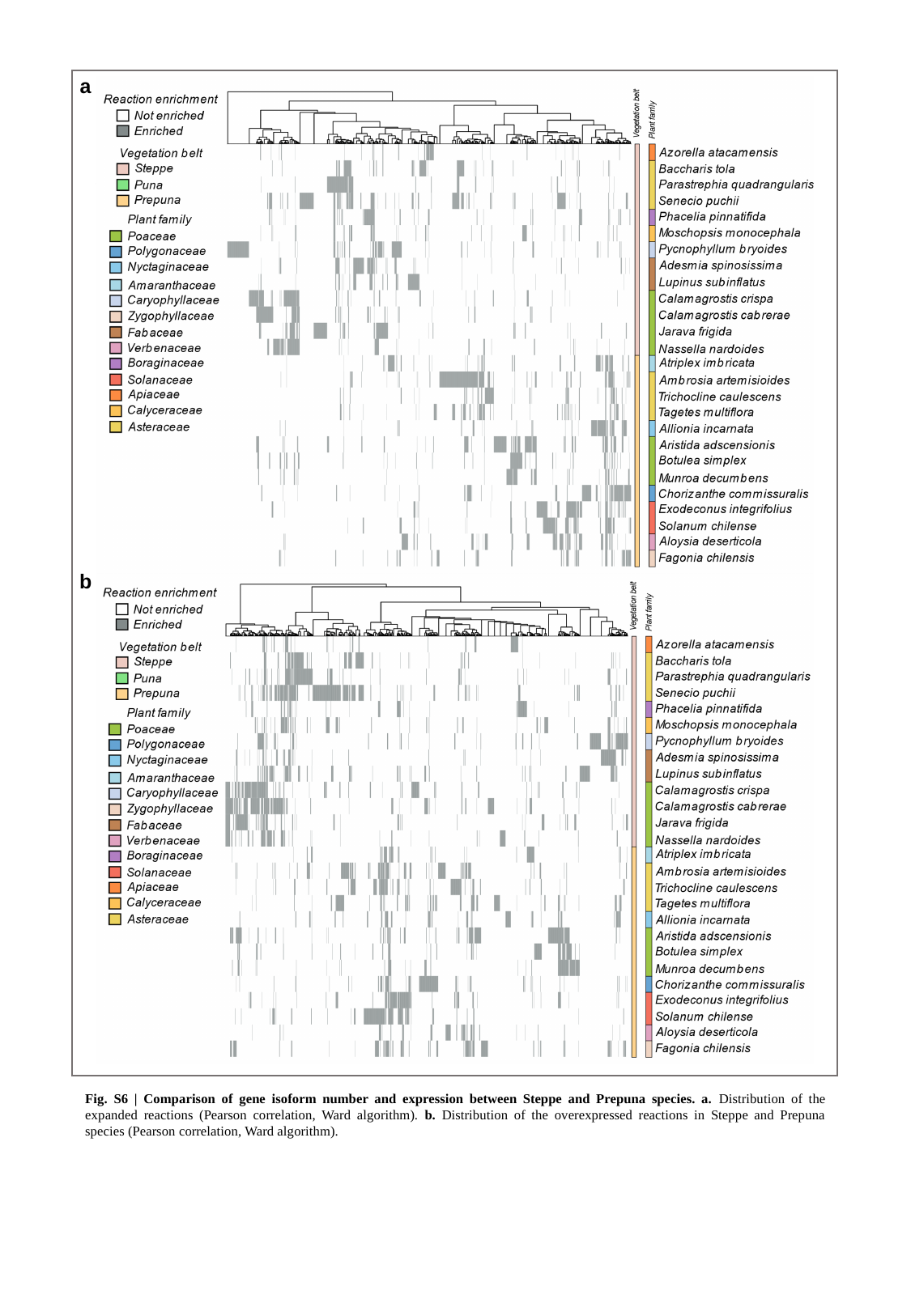

a
b
Fig. S6 | Comparison of gene isoform number and expression between Steppe and Prepuna species. a. Distribution of the expanded reactions (Pearson correlation, Ward algorithm). b. Distribution of the overexpressed reactions in Steppe and Prepuna species (Pearson correlation, Ward algorithm).
