## Supplemental Figure 7 for "Phylogenetically diverse wild plant species use common biochemical strategies to thrive in the Atacama Desert"

### Slide 1
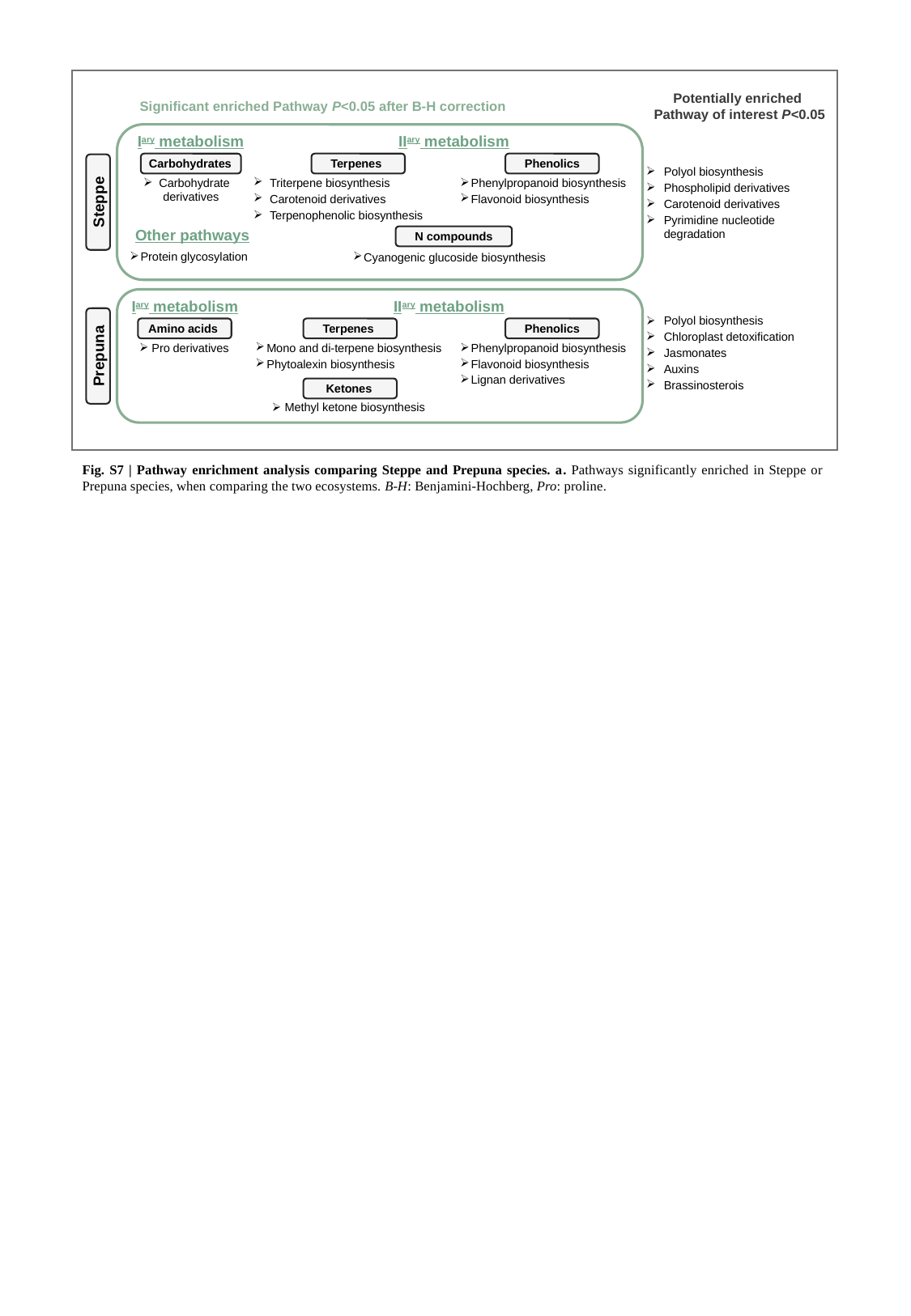

Potentially enriched
Pathway of interest P<0.05
Significant enriched Pathway P<0.05 after B-H correction
I^ary^ metabolism
Carbohydrates
 Carbohydrate derivatives
II^ary^ metabolism
Terpenes
 Triterpene biosynthesis
 Carotenoid derivatives
 Terpenophenolic biosynthesis
Phenolics
 Phenylpropanoid biosynthesis
 Flavonoid biosynthesis
N compounds
 Cyanogenic glucoside biosynthesis
 Polyol biosynthesis
 Phospholipid derivatives
 Carotenoid derivatives
 Pyrimidine nucleotide
 degradation
Steppe
Other pathways
 Protein glycosylation
I^ary^ metabolism
II^ary^ metabolism
 Polyol biosynthesis
 Chloroplast detoxification
 Jasmonates
 Auxins
 Brassinosterois
Amino acids
Terpenes
Phenolics
 Pro derivatives
 Mono and di-terpene biosynthesis
 Phytoalexin biosynthesis
 Phenylpropanoid biosynthesis
 Flavonoid biosynthesis
 Lignan derivatives
Prepuna
Ketones
 Methyl ketone biosynthesis
Fig. S7 | Pathway enrichment analysis comparing Steppe and Prepuna species. a. Pathways significantly enriched in Steppe or Prepuna species, when comparing the two ecosystems. B-H: Benjamini-Hochberg, Pro: proline.
