## Supplemental Figure 8 for "Phylogenetically diverse wild plant species use common biochemical strategies to thrive in the Atacama Desert"

### Slide 1
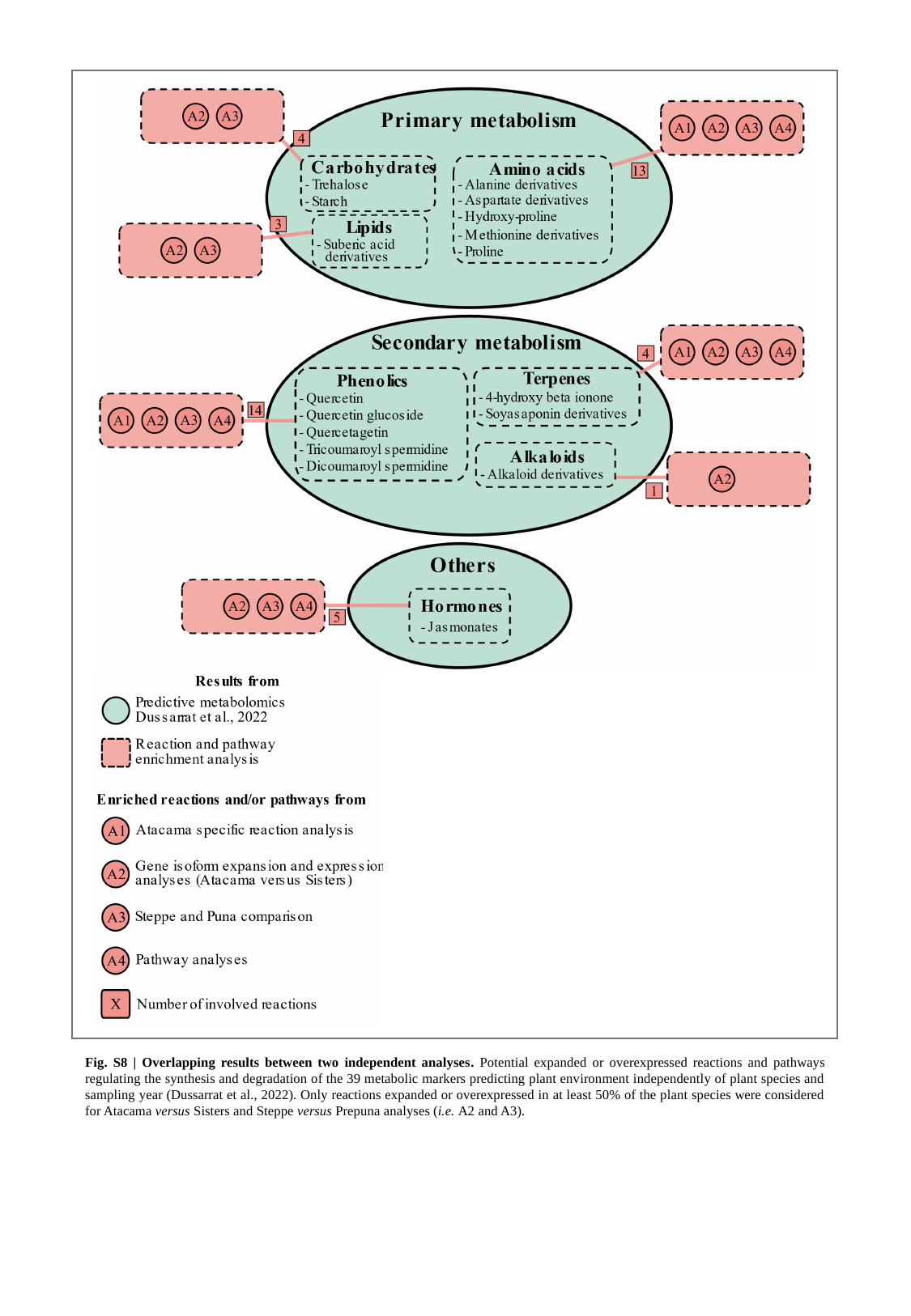

Fig. S8 | Overlapping results between two independent analyses. Potential expanded or overexpressed reactions and pathways regulating the synthesis and degradation of the 39 metabolic markers predicting plant environment independently of plant species and sampling year (Dussarrat et al., 2022). Only reactions expanded or overexpressed in at least 50% of the plant species were considered for Atacama versus Sisters and Steppe versus Prepuna analyses (i.e. A2 and A3).
