## Supplemental Figure 9 for "Phylogenetically diverse wild plant species use common biochemical strategies to thrive in the Atacama Desert"

### Slide 1
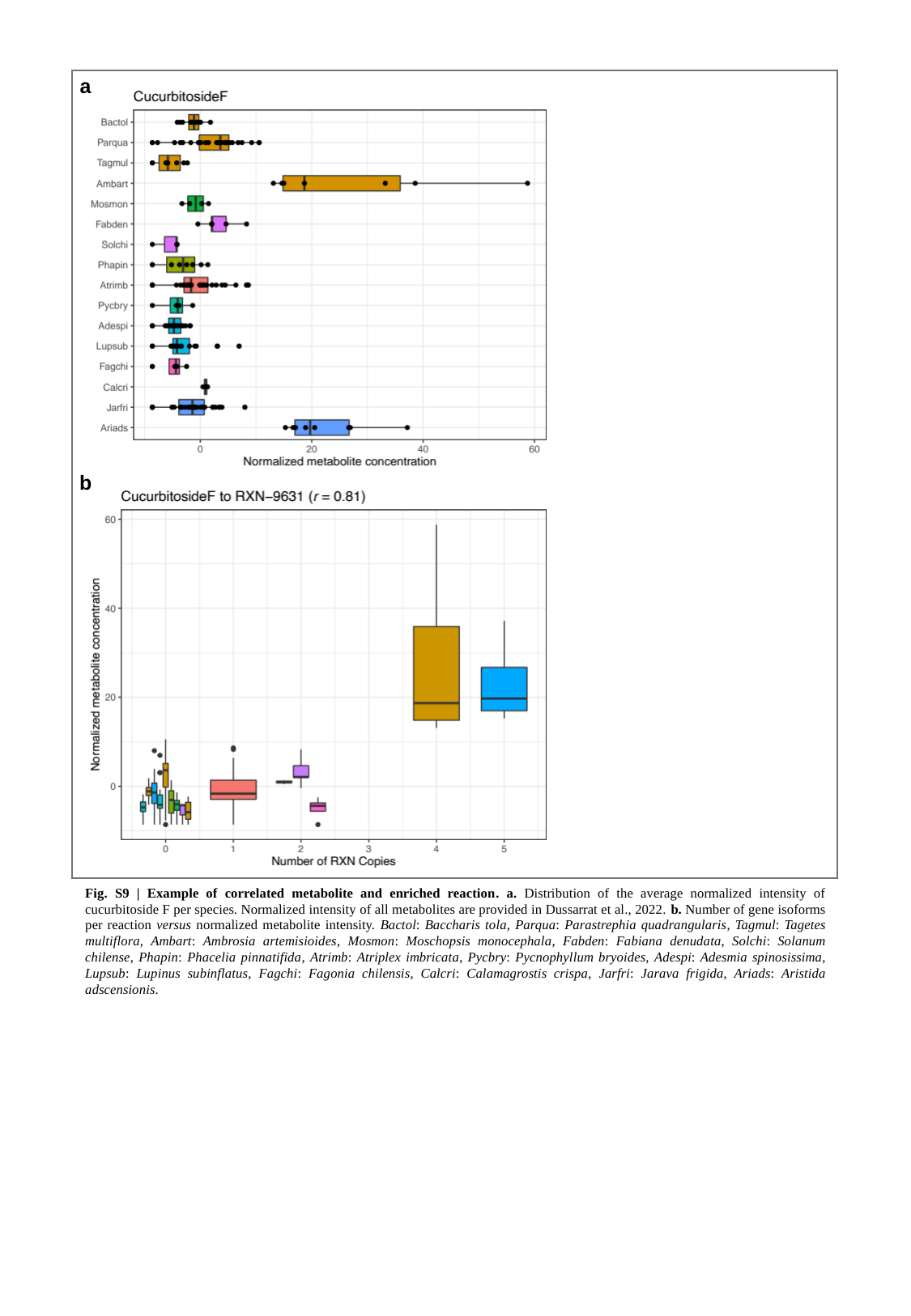

a
b
Fig. S9 | Example of correlated metabolite and enriched reaction. a. Distribution of the average normalized intensity of cucurbitoside F per species. Normalized intensity of all metabolites are provided in Dussarrat et al., 2022. b. Number of gene isoforms per reaction versus normalized metabolite intensity. Bactol: Baccharis tola, Parqua: Parastrephia quadrangularis, Tagmul: Tagetes multiflora, Ambart: Ambrosia artemisioides, Mosmon: Moschopsis monocephala, Fabden: Fabiana denudata, Solchi: Solanum chilense, Phapin: Phacelia pinnatifida, Atrimb: Atriplex imbricata, Pycbry: Pycnophyllum bryoides, Adespi: Adesmia spinosissima, Lupsub: Lupinus subinflatus, Fagchi: Fagonia chilensis, Calcri: Calamagrostis crispa, Jarfri: Jarava frigida, Ariads: Aristida adscensionis.
